# Proximity-labeling proteomics reveals remodeled interactomes and altered localization of pathogenic SHP2 variants

**DOI:** 10.1101/2025.02.26.640373

**Authors:** Anne E. van Vlimmeren, Lauren C. Tang, Ziyuan Jiang, Abhishek Iyer, Rashmi Voleti, Konstantin Krismer, Jellert T. Gaublomme, Marko Jovanovic, Neel H. Shah

**Affiliations:** Department of Chemistry, Columbia University, New York, NY 10027; Department of Biological Sciences, Columbia University, New York, NY 10027; Koch Institute for Integrative Cancer Biology, Massachusetts Institute of Technology, Cambridge, MA; Herbert Irving Comprehensive Cancer Center, Columbia University, New York, NY 10032

**Keywords:** SHP2, tyrosine phosphatase, protein-protein interaction, phosphotyrosine, mitochondria, TurboID

## Abstract

Missense mutations in *PTPN11*, which encodes the protein tyrosine phosphatase SHP2, are common in several developmental disorders and cancers. While many mutations disrupt auto-inhibition and hyperactivate SHP2, several do not enhance catalytic activity. Both activating and non-activating mutations could potentially drive pathogenic signaling by altering SHP2 interactions or localization. We employed proximity-labeling proteomics to map the interaction networks of wild-type SHP2, ten clinically-relevant mutants, and SHP2 bound to an inhibitor that stabilizes its auto-inhibited state. Our analyses revealed mutation- and inhibitor-dependent alterations in the SHP2 interactome, with several mutations also changing localization. Some mutants had increased mitochondrial localization and impacted mitochondrial function. This study provides a resource for exploring SHP2 signaling and offers new insights into the molecular basis of SHP2-driven diseases. Furthermore, this work highlights the capacity for proximity-labeling proteomics to detect missense-mutation-dependent changes in protein interactions and localization.

## Introduction

Missense mutations are a prevalent type of genetic variation, and they often drive human diseases^1,2^. Missense mutations can change protein function by perturbing active or regulatory sites, disrupting conformational stability and flexibility^3,4^, changing protein-protein interactions^5–7^, and altering subcellular localization^8,9^. Thus, the comprehensive characterization of a disease-associated missense mutation requires not only an analysis of the biochemical properties and signaling activities of that protein, but also an unbiased assessment of its broader cellular properties, including its localization and interaction network.

SHP2 is a protein tyrosine phosphatase with hundreds of documented disease-associated missense mutations^2,10^. This signaling enzyme has critical roles in diverse cell signaling pathways^11–13^, most notably as an activator of the Ras/MAPK pathway^14^. SHP2 contains a catalytic protein tyrosine phosphatase (PTP) domain and two phosphotyrosine-recognizing SH2 domains that regulate its localization and activation (**Figure 1A**)^15^. The N-SH2 domain inhibits the PTP domain when not bound to phosphoproteins (**Figure 1B**), whereas the C-SH2 primarily mediates localization^16^. SHP2 dysregulation by missense mutations leads to diverse diseases with a wide range of phenotypes^17,18,10,19–21,12^. Mutations in SHP2 underlie a large fraction of cases of Noonan Syndrome (NS) and Noonan Syndrome with Multiple Lentigines (NSML), two clinically distinct congenital disorders^22,23^. SHP2 mutations are also associated with leukemias, including juvenile myelomonocytic leukemia (JMML), acute myeloid leukemia (AML), and acute lymphoblastic leukemia (ALL), and are occasionally found in solid tumors^24,25^.

**Figure 1.**
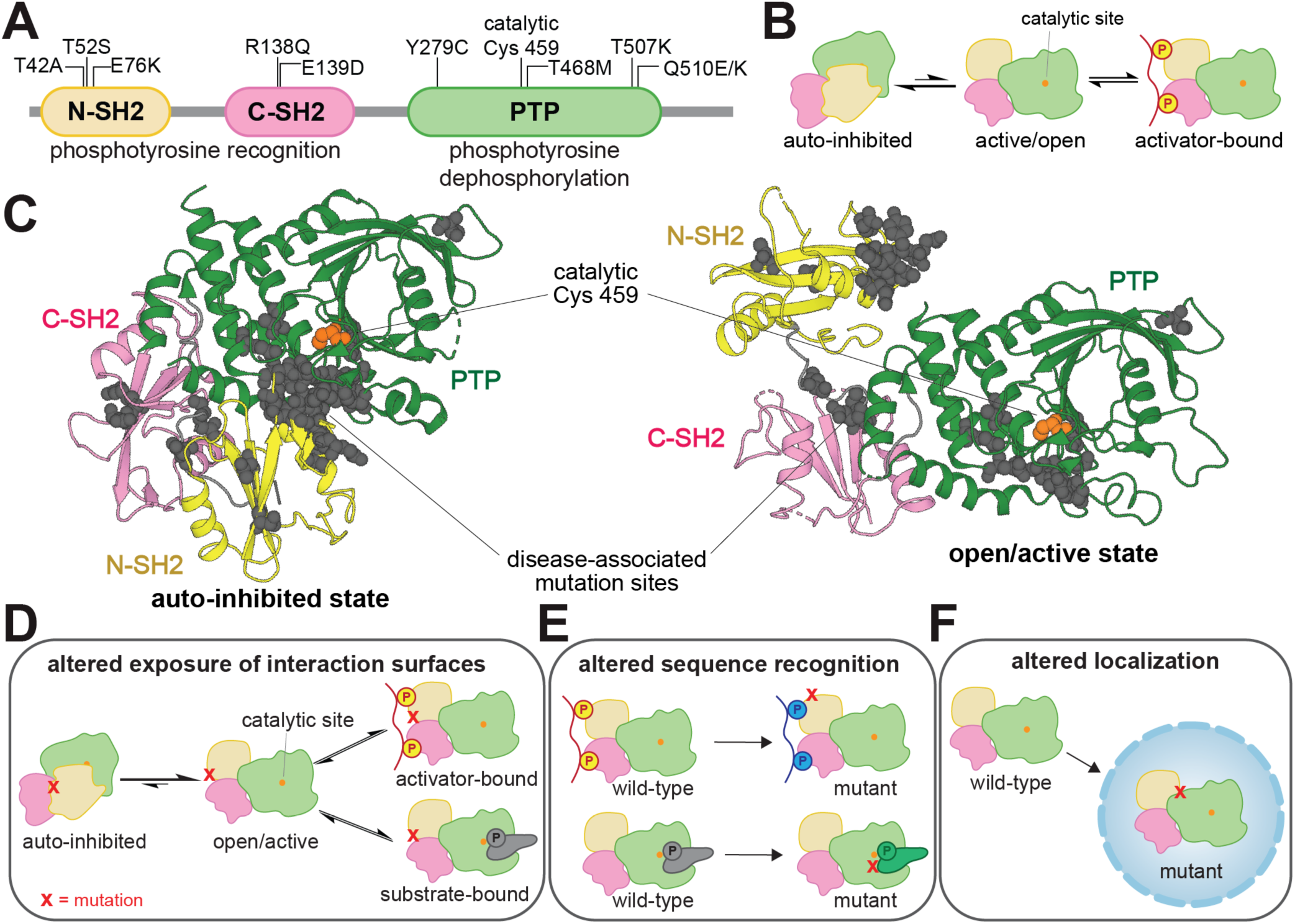
SHP2 regulation and dysregulation by mutations. (**A**) Domain architecture diagram of SHP2. Relevant mutations and the catalytic cysteine residue are labeled. (**B**) SHP2 is activated by the SH2 domains binding to phosphoproteins. (**C**) Visualization of common disease-associated mutations in SHP2 (gray spheres) in the auto-inhibited state (*left*, PDB code 4DGP) and open state (*right*, PDB code 6CRF). Cancer mutations cluster at the auto-inhibitory interface between the N-SH2 and PTP domains. Schematic diagrams showing that (**D**) mutations can alter the exposure of catalytic or non-catalytic interaction surfaces in SHP2, (**E**) mutations can alter sequence recognition in either the SH2 domain binding pockets or PTP domain active site, and (**F**) mutations can alter subcellular localization.

Most well-characterized pathogenic SHP2 mutations hyperactivate the enzyme by disrupting the auto-inhibitory interface between the N-SH2 and PTP domains (**Figure 1C**). These mutations can also alter protein-protein interactions, not only by allowing access the catalytic site, but also by putting the SH2 domains in a binding-competent state and exposing other cryptic binding interfaces^26–29^ (**Figure 1D**). Many clinically-observed mutations occur outside this interface and do not enhance basal catalytic activity^30^, suggesting alternative disease mechanisms (**Figure 1E,F**)^7,31^. For example, the T42A mutation alters the N-SH2 ligand binding pocket, changing binding affinity and specificity for phosphoproteins^7,32^. Several studies have suggested that mutations in SHP2 can rewire its role in signaling networks by remodeling protein-protein interactions^7,33,34^, underscoring the need for a broad and unbiased functional analysis of SHP2 missense mutations.

Here, we profile mutations-specific changes to the SHP2 interaction network using proximity-labeling proteomics. We report the interactomes of wild-type SHP2 (SHP2^WT^) and 10 disease-associated mutants, chosen for their diverse molecular and clinical effects. Our experiments identify mutation-driven changes in SHP2 protein-protein interactions. Our results also pinpoint a role for SHP2 in the mitochondria, as well as enhanced mitochondrial localization and interactions for select mutants. We find that destabilizing SHP2 mutations increase localization to the mitochondria, potentially affecting mitochondrial homeostasis. We also examine how the clinically-relevant allosteric inhibitor TNO155 remodels the SHP2 interactome. Our datasets expand the mechanistic paradigm for mutational effects in SHP2 and will be a resource for understanding SHP2 signaling and pathogenicity. Furthermore, our work demonstrates a robust strategy for identifying mutation-driven effects to protein function using TurboID-proximity labeling.

## Results

### Proximity-labeling proteomics identifies novel wild-type SHP2 interactors

The overarching goal of this study is to examine how mutations alter SHP2 protein-protein interactions, however we lack a comprehensive SHP2^WT^ interactome dataset. Thus, we conducted affinity-purification mass spectrometry of myc-tagged SHP2^WT^ in SHP2-knock-out HEK 293 cells to identify interacting proteins. Consistent with previous work^35^, this experiment identified few interacting proteins, none of which were known SHP2 interactors (**Supplementary Figure 1A and Supplementary Table 1**). Furthermore, SHP2^T42A^, a mutant with enhanced N-SH2 binding affinity did not pull down more phosphoproteins (**Supplementary Figure 1B and Supplementary Table 1**). Signaling processes often require transient protein-protein interactions in order to be dynamic and reversible, and this may preclude facile identification of interactors by immunopurification^36^. Indeed, PTP and SH2 domains are known to have weak and transient interactions with phosphoproteins, making both substrate and interactor identification by traditional immunopurification methods difficult^37,38^. Given these innate features of signaling proteins, coupled with our low-yielding affinity-purification mass spectrometry data, we reasoned that an alternative method was needed to map the SHP2 interactome.

To identify transient SHP2 interactions, co-localized proteins, and mutational differences, we implemented proximity-labeling proteomics using the engineered high-activity biotin ligase TurboID (**Figure 2A**)^39^. Previous studies have used the low-activity biotin ligase BirA to analyze or validate the interactomes of various tyrosine phosphatases^40–42^, however, the data for SHP2 are sparse. We fused TurboID to the C-terminus of SHP2, to preserve N-SH2 mediated auto-inhibition. We confirmed that SHP2^WT^-TurboID retains catalytic activity *in vitro* and can be activated by SH2-domain engagement of phosphopeptides (**Supplementary Figure 2A**). We also confirmed the hyperactivating effect of the E76K (**Supplementary Figure 2B**). Furthermore, we ensured that SHP2-TurboID retains activity in cells and that this activity can be further enhanced by the E76K mutation (**Supplementary Figure 2C-E**).

**Figure 2.**
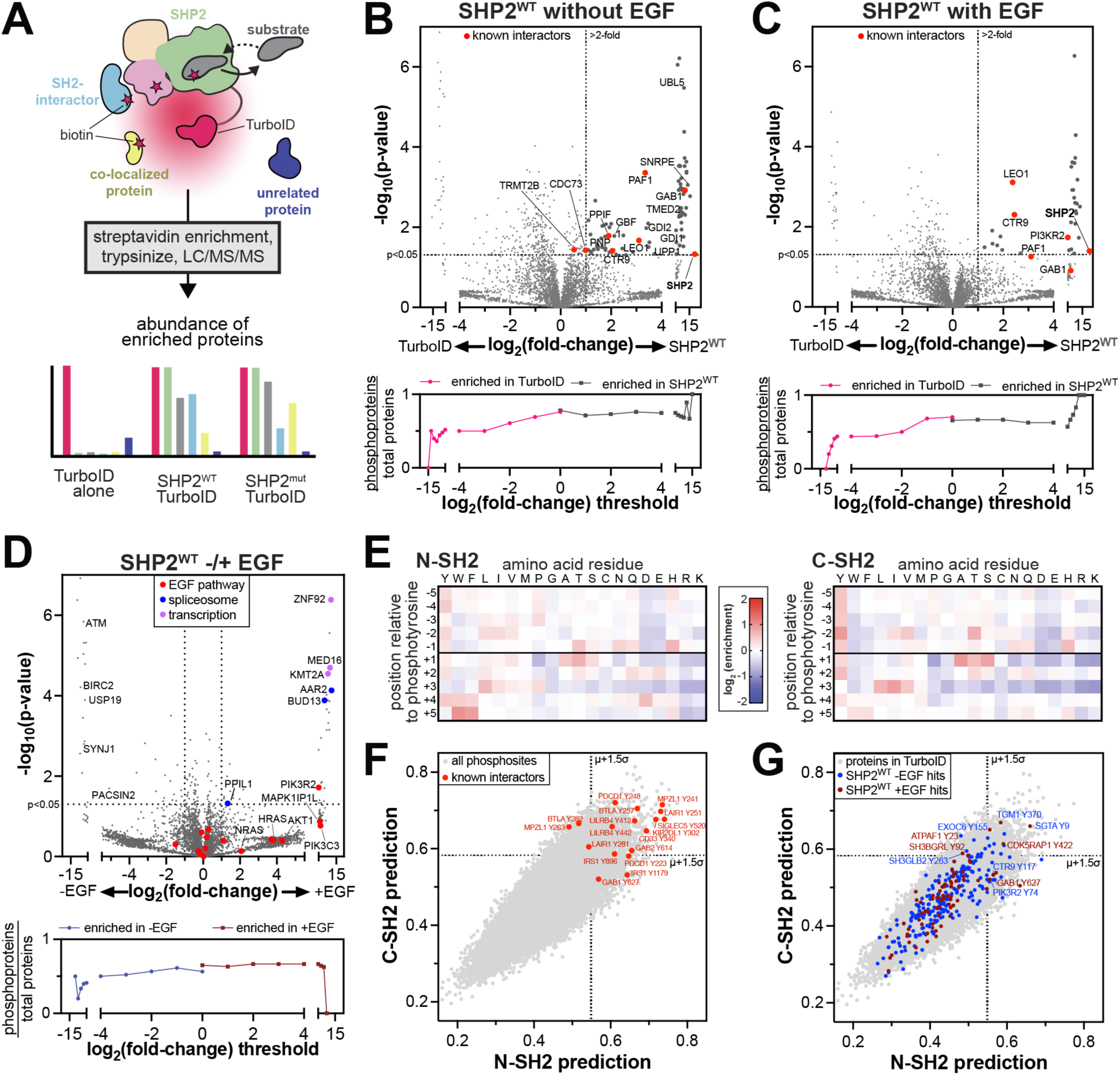
Profiling the interactomes of wild-type SHP2. (**A**) Schematic diagram of SHP2-TurboID experiments. TurboID generates reactive biotin derivatives, which label substrates, interactors, and co-localized proteins. MS/MS can be used to detect and quantify the interactome. (**B**) *top:* Comparison of SHP2^WT^-TurboID with a TurboID-only control, in absence of EGF stimulation. Several known SHP2 interactors were identified, including proteins that are completely absent from the TurboID control. Gaps in the x-axis exclude non-significant points. Points at extreme x-values reflect proteins that were abundant in all SHP2-TurboID replicates and completely absent (imputed) in the TurboID-only control, or vice versa. *Bottom*: Analysis of the total number of potential phosphoproteins, as identified by PhosphoSitePlus, per total number of observed proteins above each fold-change threshold. (**C**) Same as (**B**) but with EGF stimulation. (**D**) *top*: Comparison of SHP2^WT^-TurboID in unstimulated and EGF-stimulated conditions. Proteins involved in trascription, spliceosome, and the EGF pathway are indicated. *Bottom*: Analysis of the total number of potential phosphoproteins, as identified by PhosphoSitePlus, per total number of observed proteins above each fold-change threshold, for SHP2^WT^-TurboID with and without EGF stimulation. (**E**) Heatmaps depicting position-specific weight matrices for peptide recognition by the SHP2 N-SH2 and C-SH2 domains. (**F**) Predicted scores for binding to the N-SH2 and C-SH2 domains for all phosphotyrosine sites in PhosphoSitePlus (gray dots). Phosphosites on several known interactors are labeled (red dots). (**G**) Predicted scores for binding to the N-SH2 and C-SH2 domains for all phosphosites on proteins in our proteomics dataset (gray dots). Phosphosites on proteins in the SHP2^WT^ interactomes are colored in blue (no EGF) and dark red (+EGF).

We next expressed SHP2^WT^-TurboID and a TurboID-only control in SHP2 knock-out HEK 293 cells to profile the SHP2^WT^ interactome^43^. SHP2-TurboID was expressed at comparable levels to endogenous SHP2 in wild-type HEK 293 cells, however we observed partial cleavage of SHP2 from TurboID (**Supplementary Figure 2F**). Following established protocols, biotinylated proteins were enriched and identified by mass spectrometry^44^. We identified 78 proteins with significant enrichment for SHP2^WT^ over the TurboID-only control, applying a p-value cut-off of <0.05 and filtering for proteins with a log_2_ fold-change of >1 (**Figure 2B**, *top*, **and Supplementary Table 2**). Among the significantly enriched proteins were known SHP2 interactors, such as components of the Paf1 complex (Paf1, Ctr9, Leo1) and the scaffold protein Gab1 (**Figure 2B**, *top*)^45–48^ We also observed enrichment of the mitochondrial peptidyl-prolyl isomerase PPIF, a protein previously shown to co-fractionate with SHP2, giving further confidence to the identified interactome^49^.

In addition to known interactors, our data show that SHP2 interacts or co-localizes with components of several functionally diverse protein complexes and pathways **(Supplementary Figure 3A and Supplementary Table 2)**. For example, we identified GDI1 and GDI2, which regulate Rab proteins; and UBL5 and SNRPE, which are components of the spliceosome. Previously, SNRNP27, another spliceosome component was identified as a putative interactor for SHP2^50^. We also identified GBF1 and TMED2, which coordinate Golgi trafficking of COPI vesicles; as well as UPP1 and PNP, enzymes involved in pyrimidine and purine metabolism, respectively. Finally, we observed a distinctive signal for many mitochondrial proteins, including components of the electron transport chain, which we discuss extensively below. To confirm that these results are not due to increased protein abundance in SHP2-expressing cells, we also analyzed the total proteome of the matched cell lysates from which our TurboID experiments were performed. Proteins that were significantly enriched in the SHP2^WT^-TurboID experiments did not show substantial changes in protein abundance (**Supplementary Figure 2G and Supplementary Table 3**).

Next, we stimulated the epidermal growth factor receptor (EGFR), which phosphorylates many downstream proteins that bind to and activate SHP2^48^. We compared the interactome of SHP2^WT^-TurboID to the TurboID-only control in cells stimulated with 100 ng/mL EGF and identified 31 proteins that were significantly enriched by SHP2^WT^ (**Figure 2C**, *top*, **Supplementary Figure 3B, and Supplementary Table 2**). A few of these proteins, including Gab1 and components of the Paf1 complex, were also observed in unstimulated samples. However, most proteins were uniquely enriched in the EGF-stimulated samples. For example, we detected PI3KR2, a regulatory subunit of phosphatidylinositol-3 (PI3) kinase, which is a component of the EGF pathway and an established SHP2 interactor^51^. As done for the unstimulated sample, we confirmed that enriched proteins were not the result of increased protein abundance (**Supplementary Figure 2H and Supplementary Table 3**). Direct comparison of SHP2^WT^-TurboID with and without EGF stimulation revealed 25 proteins with EGF-dependent enhanced proximity-labeling, including several components of the EGF pathway, along with proteins involved in transcription and RNA splicing (**Figure 2D**, *top*). Interestingly, 30 proteins showed reduced biotinylation upon EGF stimulation, including PACSIN2 and SYNJ1, which are involved in EGFR endocytosis in absence of EGF stimulation^52,53^. We also identified BIRC2, a modulator of mitogenic kinase signaling, and its interactor USP19^54,55^, as well as checkpoint kinase ATM. Collectively, these experiments reveal the SHP2^WT^ interactome, its restructuring by EGF stimulation, and several potentially novel functions for SHP2. Protein tyrosine phosphorylation partly shapes the SHP2^WT^ interactome

Since all three globular domains of SHP2 engage tyrosine-phosphorylated proteins, we reasoned that a fraction of the SHP2 interactome would depend on tyrosine phosphorylation, particularly in cells with increased abundance of phosphoproteins due to EGF stimulation. To test this hypothesis, we used the PhosphoSitePlus database to assess what fraction of the proteins enriched in our TurboID dataset have potential tyrosine phosphorylation sites^56^. Proteins enriched by SHP2^WT^-TurboID were more likely to have tyrosine phosphorylation sites than those enriched by the control (**Figure 2B,C***, bottom*). Furthermore, we found that the EGF-stimulated samples had a slight bias for proteins with tyrosine phosphorylation sites when compared with unstimulated samples (**Figure 2D***, bottom*). This analysis demonstrates that SHP2 interactions in our datasets are partly driven by tyrosine phosphorylation.

Next, we used the sequence specificities of the N- and C-SH2 domains to identify potential SH2 domain binding sites in our TurboID datasets. We conducted a high-throughput binding assay with the SHP2 N- and C-SH2 domains, using a degenerate ∼10^6^ peptide library containing random sequences with a central phosphotyrosine (**Figure 2E and Supplementary Table 4**)^57^. This approach yielded position-specific weight matrices, which we used to score the ∼40,000 tyrosine phosphorylation documented in the PhosphoSitePlus database (**Supplementary Table 4**)^56,57^. We confidently identified several known SHP2 binding sites as high-affinity sequences (**Figure 2F**). In the SHP2^WT^-TurboID interactome, we identified 26 and 10 predicted tight binding phosphosites in unstimulated and EGF-stimulated conditions, respectively (**Figure 2G**). These include phosphosites on known SHP2 binders, such as Gab1, Ctr9, and PI3KR2, as well as potential novel interactors, such as EXOC6, which has been linked to both EGFR and PI3 kinase signaling^58^, and SH3BGRL proteins, which modulate phosphotyrosine signaling through direct interaction with EGFR-family kinases^59^. This juxtaposition of biochemical and proteomic datasets provides additional mechanistic insights into specific SHP2 interactions.

### Pathogenic mutations have divergent effects on the SHP2 interactome

Having established a robust method to map the SHP2^WT^ interactome, we then examined mutation-dependent changes in protein-protein interactions. We selected ten disease-associated SHP2 mutants, chosen for their distribution across the protein, varying effects on structure and activity, and diverse clinical outcomes (**Figure 3A**). Some of these mutants have been thoroughly characterized whereas others remain more elusive. The Noonan Syndrome-associated T42A mutation enhances N-SH2 domain phosphoprotein binding affinity and specificity with a marginal effect on basal phosphatase activity^7,32,60^. The T52S mutation, identified in JMML patients, does not alter basal activity, ligand-binding affinity, or ligand-binding specificity^7^, and its pathogenic mechanism is unknown. E76K is a well-studied JMML mutation that disrupts auto-inhibition, yielding a constitutively active state of SHP2^61,62^. The melanoma mutation R138Q severely attenuates binding of the C-SH2 domain to phosphoproteins and might adopt a slightly more open protein conformation^7^. Curiously, the NS and JMML mutation E139D causes a substantial increase in activity despite not being at the auto-inhibitory interface, and reports of its effect on ligand-recognition by the C-SH2 domain have been conflicting^7,32^. Y279C and T468M, both located in the PTP domain and found in NSML, reduce SHP2 catalytic efficiency while destabilizing the auto-inhibited state^63^. T507K has been identified in solid tumors and alters substrate specificity^34^. Finally, Q510K and Q510E both reduce catalytic activity^30^, however Q510E has been identified in NSML and ALL, whereas Q510K has only been reported in patients with ALL. The interactomes and localization of most of these mutants have not been systematically explored.

**Figure 3.**
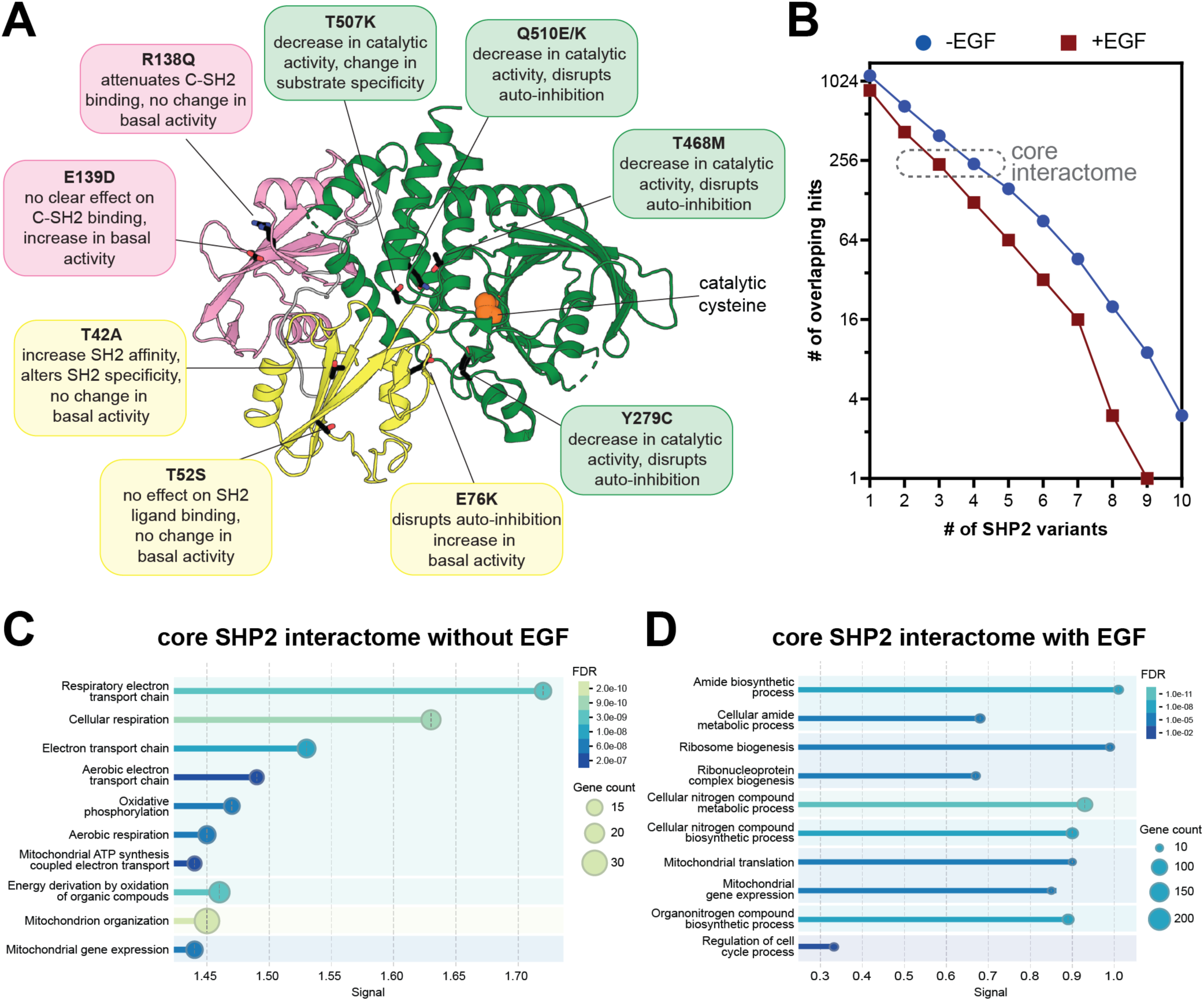
The interactomes of diverse pathogenic SHP2 variants. (**A**) Overview of mutations in this study. Their position on the auto-inhibited structure (PDB code 4DGP) are indicated, as are known characteristics of each mutant. (**B**) Number of overlapping proteins that are significantly enriched over the negative control for multiple SHP2 variants, both with and without EGF stimulation. (**C**) GO enrichment analysis for biological processes of the core SHP2 interactome without EGF stimulation. (**D**) GO enrichment analysis for biological processes of the core SHP2 interactome with EGF stimulation.

We conducted TurboID proximity labeling with each of these mutants analogous to our SHP2^WT^ experiments. Interactomes for each mutant were defined by comparison with the TurboID-only control, using the same p-value (<0.05) and enrichment cut-off (>2-fold) used for SHP2^WT^ (**Supplementary Table 2**). In the absence of EGF stimulation, the SHP2 mutants shared between 13 and 39 hits with SHP2^WT^, which showed enrichment for 78 proteins over the TurboID-control (**Supplementary Figure 4A**). In the presence of EGF, the interactome of each mutant was generally smaller (**Supplementary Figure 4B**). One plausible explanation for this is that, in unstimulated cells, SHP2 is more diffusely localized throughout the cell, whereas in EGF-stimulated cells, membrane-proximal tyrosine kinase activity leads to recruitment of SHP2 to specific subcellular locations. To accommodate potential data sparsity, we also calculated a relaxed pairwise overlap, where both p-value and fold-change cut-offs were applied to the first variant, but only one of these cut-offs was applied to the second variant (**Supplementary Figure 4C**,D). This revealed substantial overlap between SHP2 variants (e.g. between 52 and 61 shared proteins for each mutant with SHP2^WT^ out of the 78 hits without EGF stimulation), providing further confidence in the interactors we identified.

Using the strict combined p-value and enrichment cut-offs, we defined a core SHP2 interactome as proteins significantly enriched over the TurboID-only control for at least 4 out of the 11 SHP2 variants in unstimulated cells and 3 out of 11 variants with EGF stimulation, approximately 240 proteins in each case (**Figure 3B**). Gene ontology analysis on this core interactome revealed conserved mitochondrial localization and interactions across SHP2 mutants (**Figure 3C,D and Supplementary Figure 5**). Both with and without EGF stimulation, these core SHP2 interactomes included mitochondrial ribosome proteins, but components of the respiratory chain complexes (e.g. NDUF and COX proteins) were distinctively enriched in the absence of EGF. By contrast, many cell cycle regulatory proteins were enriched in the EGF-stimulated core SHP2 interactome (e.g. CDK4 and CDC27), as was the EGFR-family kinase Her2 (ERBB2).

Beyond this core interactome, individual mutants displayed distinct interactomes from SHP2^WT^ and from each other (**Supplementary** Figures 4A-D **and Supplementary Table 2**). While some of these differences may arise from data sparsity or noise in our datasets, we hypothesize that many differences reflect mutation-induced changes in protein conformation or molecular recognition, as discussed in subsequent sections. Collectively, the analyses in this section show that pathogenic SHP2 mutants generally preserve some core functions of wild-type SHP2, but specific mutations partly remodel the SHP2 interactome, and this can be further altered by EGF stimulation.

### Mutation-dependent changes in phosphoprotein recognition underlie altered SHP2 interactomes

Since many SHP2 interactions are driven by phosphoprotein binding, we first focused on the T42A and R138Q mutations in the N- and C-SH2 domains, which are known to impact phosphoprotein recognition (**Figure 3A**). We and others previously showed that the T42A mutation enhances the affinity and alters specificity of the N-SH2 domain for phosphoproteins^7,32^. Consistent with this we observed an overall increase in the number of proteins enriched by SHP2^T42A^ over SHP2^WT^ in our proximity-labeling datasets (**Figure 4A and Supplementary Figure 4A-D**). Among these SHP2^T42A^-enriched proteins were MPZL1 and ACOX1 (**Figure 4B**). In our previous work using phosphopeptide libraries to profile SHP2 SH2 mutants, we found that phosphopeptides derived from these proteins showed enhanced binding to N-SH2^T42A^ with respect to N-SH2^WT^, providing a mechanistic explanation for the enrichment of these proteins in our SHP2^T42A^ interactome (**Figure 4C**)^7^.

**Figure 4.**
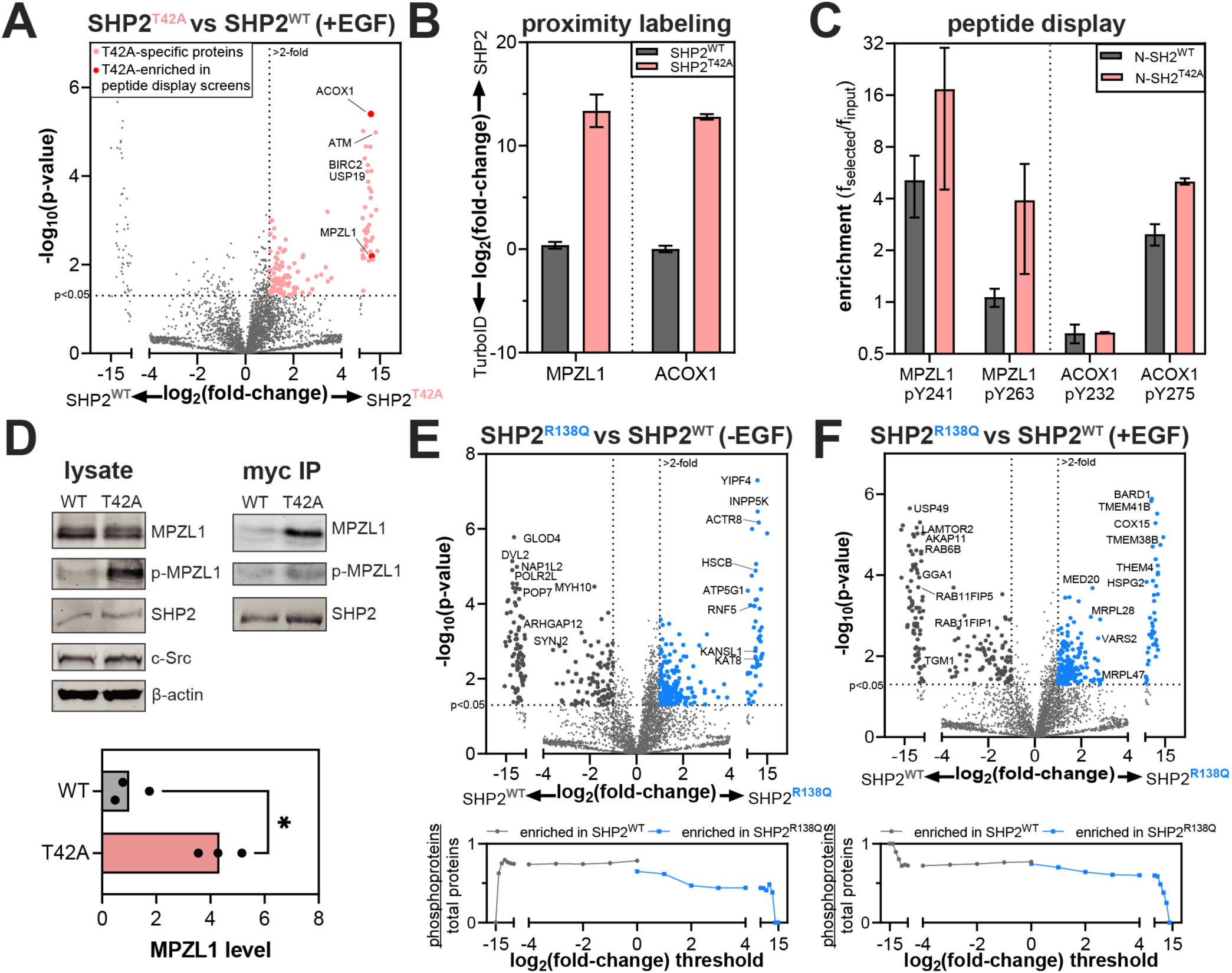
Rewired interactomes in SHP2 mutants with altered phosphoprotein binding properties. (**A**) Volcano plot showing enriched proteins in SHP2^T42A^, compared to SHP2^WT^. (**B**) Log_2_ fold-change in enrichment for MPZL1 and ACOX1 by SHP2^WT^ and SHP2^T42A^ compared with the TurboID control. (**C**) Enrichment of MPZL1- and ACOX1-derived phosphopeptides from peptide display screens using the SHP2 N-SH2^WT^ and N-SH2^T42A^ domains. (**D**) Phosphorylation levels and co-immunopurification of MPZL1 by SHP2^WT^ and SHP2^T42A^. Quantified MPZL1 levels in the myc-SHP2 co-IP are shown in the graph. Asterisk denotes p<0.05. (**E**) (*top*) Volcano plots highlighting enriched proteins in SHP2^R138Q^, compared to SHP2^WT^ in the absence of EGF stimulation. (*bottom)* Analysis of the total number of potential phosphoproteins, as identified by PhosphoSitePlus, per total number of observed proteins past each fold-change threshold for SHP2^WT^ versus SHP2^R138Q^. The SHP2^R138Q^ interactome is less enriched in potentially tyrosine-phosphorylated proteins than the SHP2^WT^ interactome. No negative selection against tyrosine phosphorylation sites is observed for SHP2^WT^. (**F**) Same as (**E**), but with EGF stimulation.

MPZL1 is a known SHP2 interactor, and enhancement of this interaction through SHP2 NSML mutations is linked to hypertrophic cardiomyopathy^28^. Unlike SHP2^T42A^, which is a Noonan Syndrome mutant, NSML mutants like SHP2^T468M^ and SHP2^Q510E^ have a higher propensity to adopt the open conformation, allowing them to more readily bind phosphoproteins, and these mutants generally also have impaired catalytic activity^31^. In NSML, MPZL1 is hyperphosphorylated on Y241 and Y263, because open conformation SHP2 mutants are more competent for MPZL1 binding and protect these sites from dephosphorylation^64^. Indeed, like SHP2^T42A^, the NSML SHP2^T468M^ and SHP2^Q510E^ mutants showed increased interaction with MPZL1 in our TurboID data relative to SHP2^WT^ (**Supplementary Figure 4E**). We validated the enrichment of MPZL1 by SHP2^T42A^ by co-immunopurification, which showed that the T42A mutation both enhances MPZL1 binding and shows increased in MPZL1 phosphorylation (**Figure 4D**). This demonstrates that SHP2^T42A^, despite not adopting a more open conformation than SHP2^WT^, converges on increased interaction with MPZL1 through altered sequence-recognition in the N-SH2 domain^7^.

The R138Q mutation severely weakens phosphotyrosine binding to the C-SH2 domain by removing a conserved Arg residue^7^. Surprisingly, in our proximity-labeling experiments we observed more proteins enriched with SHP2^R138Q^ than SHP2^WT^, suggestive of increased interactions or co-localization with other proteins (**Figure 4E,F**, *top***, and Supplementary Figure 4A-D**). Since this mutation disrupts phosphoprotein interactions, we suspected that proximity-labeling with SHP2^R138Q^ would be less dependent on tyrosine-phosphorylated proteins than with SHP2^WT^. Indeed, proteins enriched by SHP2^R138Q^ are less likely to be phosphoproteins than those enriched by SHP2^WT^ (**Figure 4E,F**, *bottom*)^56^. Since the C-SH2 domain of SHP2 is critical for localization, one plausible explanation for the increased proximity-labeling with SHP2^R138Q^ is that the loss of C-SH2 function leads to SHP2 mislocalization and promiscuous protein-protein interactions. Overall, our analyses of SHP2^T42A^ and SHP2^R138Q^ reveal how changes in phosphoprotein recognition can measurably alter the interactome of a signaling protein.

### Pathogenic mutations remodel the SHP2 mitochondrial interactome

As noted above, mitochondrial proteins constitute a large part of the core SHP2 interactome (**Figure 3C,D and Supplementary Figure 5**). SHP2^WT^ showed enrichment of 21 mitochondrial proteins over the TurboID-only control in unstimulated cells (**Figure 5A**), consistent with previous work demonstrating mitochondrial localization of SHP2^65–71^. We further validated the localization of endogenous SHP2 to the mitochondria by fluorescence microscopy (**Supplementary Figure 6A**). Levels of biotin in the mitochondria differ from those in the cytoplasm or other organelles, which could bias subcellular TurboID signal^72–74^. In the absence of exogenous biotin, we found no bias in mitochondrial biotinylation, and the distinctive SHP2-dependent mitochondrial signal was only measurable when exogenous biotin was added (**Supplementary Figure 6B**). Notably, proximity-labeling of mitochondrial proteins was stronger in unstimulated cells compared with EGF-stimulated cells, suggesting that the mitochondrial localization of SHP2 is dependent on its signaling state (**Supplementary Figure 6B**).

**Figure 5.**
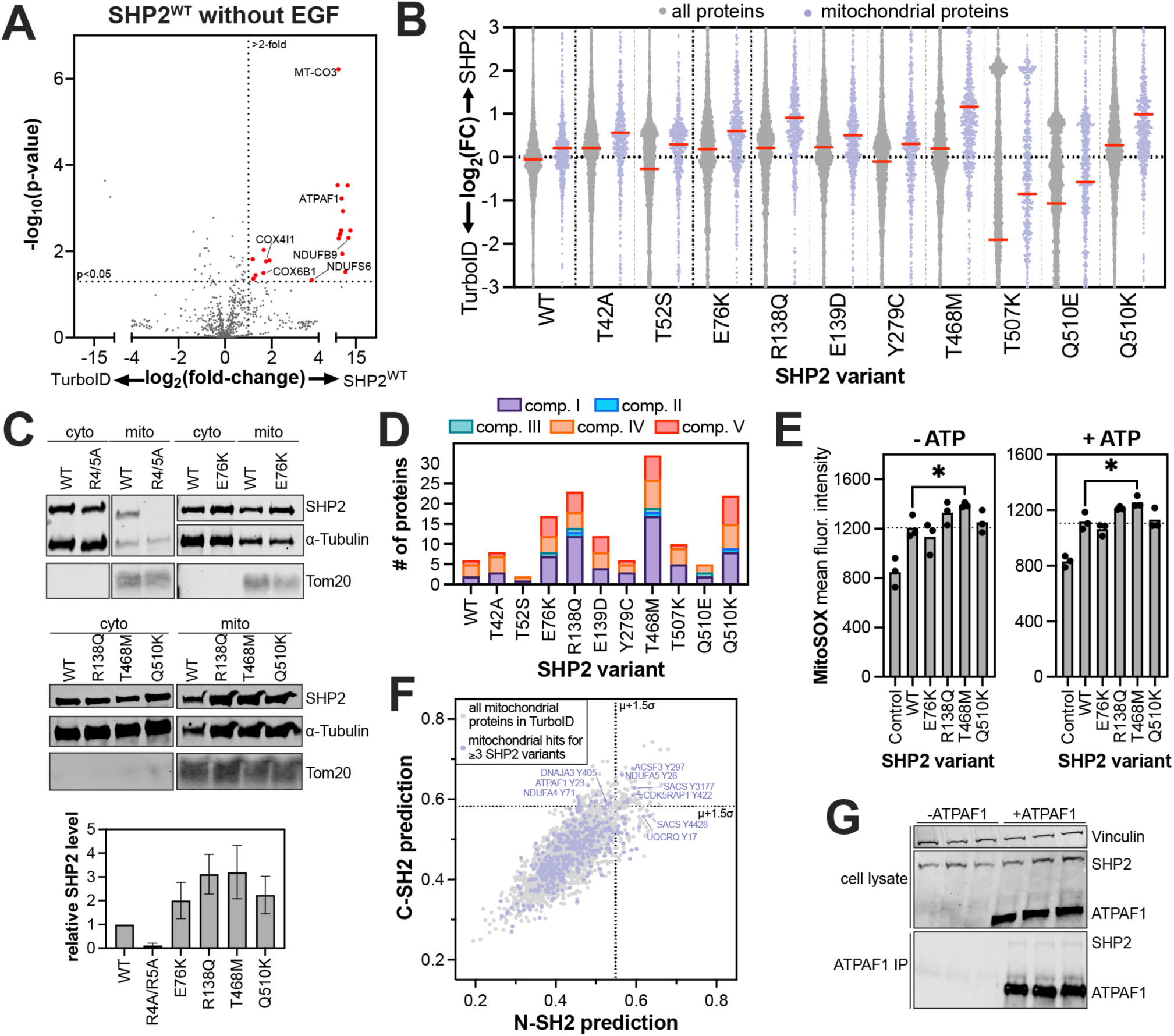
SHP2 interactions in the mitochondria. (**A**) Volcano plot of the mitochondrial proteins in our TurboID dataset, showing many mitochondrial proteins that are significantly enriched by SHP2^WT^-TurboID compared to the TurboID-only control, in absence of EGF stimulation (red dots). (**B**) Enhanced proximity labeling of mitochondrial proteins by SHP2 variants relative to the TurboID-only control (lilac), compared to proximity labeling of the whole proteome (gray). (**C**) Cellular fractionation and western blots confirming enhanced mitochondrial localization of SHP2^E76K^, SHP2^R138Q^, SHP2^T468M^, and SHP2^Q510K^, relative to SHP2^WT^. SHP2^R4/5A^ (a R4A+R5A double mutant), which has a defective mitochondrial localization signal, was used as a control and shows reduced mitochondrial abundance. (**D**) The number of electron transport chain proteins significantly enriched over TurboID for each SHP2 variant in the absence of EGF stimulation. (**E**) MitoSOX staining showing increased reactive oxygen species for SHP2^T468M^ (-/+ ATP). Asterisk denotes p<0.05. (**F**) N-SH2 and C-SH2 predictions for human phosphosites. Select high-scoring mitochondrial phosphosites are labeled. (**G**) Co-immunopurification of SHP2^WT^ by ATPAF1, co-expressed in HEK293 cells. Three independent replicates are shown on the same blots.

Interestingly, we noticed a distinctive increase in mitochondrial signal for R138Q, T468M, Q510K, and to a lesser extent, E76K (**Figure 5B and Supplementary Figure 7**). To validate this, we expressed these mutants without TurboID fusion in SHP2 knock-out HEK 293 cells, isolated mitochondria, blotted for SHP2, and confirmed that some SHP2 mutants have more mitochondrial localization than SHP2^WT^ (**Figure 5C**). SHP2-associated Noonan Syndrome and cardiomyopathies have symptoms that resemble those in primary mitochondrial disorders^75^, and disease-associated activating SHP2 mutants have been shown to impact mitochondrial function and homeostasis^71,76^. Thus, the increased mitochondrial localization for some mutants could contribute to pathogenicity, warranting a deeper analysis of the specific interactions and functions of SHP2 in this organelle.

SHP2 has been reported to interact with proteins in complex I and III of the electron transport chain, and SHP2 mutations can modulate mitochondrial respiration^71,77^. Consistent with this, our data show that many SHP2 variants have enhanced proximity labeling of electron transport chain proteins relative to the TurboID-only control (**Figure 5D**). SHP2^WT^ primarily labeled components of complex I and complex IV, while some mutants showed different labeling patterns. For example, whereas SHP2^WT^ only labeled accessory subunits of complex I, SHP2^T468M^ also labeled assembly- and core-subunits, suggestive of a longer residency near complex I. Notably, we saw no labeling of complex III by SHP2^WT^, and this was recapitulated in most SHP2 mutants. By contrast, several mutants showed an increase in labeling of complex V (ATP synthease) proteins (**Figure 5D**). These data suggest that SHP2 mutants could potentially alter core mitochondrial functions, such as oxidative phosphorylation and ATP production. Indeed, we found that a few SHP2 mutants, particularly SHP2^T468M^, enhanced formation of mitochondrial reactive oxygen species, indicating mutant-specific effects on respiration (**Figure 5E**).

Finally, we sought to identify new binding sites for SHP2 in the mitochondria that could inform future functional investigations. There were 165 mitochondrial proteins enriched by at least three SHP2 variants (either with or without EGF stimulation). We scored the 400 putative tyrosine phosphorylation sites on these proteins, as described above (**Figure 2E-G**)^57^. We found high-scoring phosphosites on NDUFA5, UQCRQ, and NDUFA4, components of complex I, III, and IV of the electron transport chain, respectively (**Figure 5F and Supplementary Table 4**). We also identified a high-scoring phosphosite on ATPAF1, an assembly factor of the F(1) component of ATP synthase, and validated that SHP2 can directly interact with ATPAF1 via co-immunopurification (**Figure 5G**). Previously, SHP2 has also been shown to directly bind to NDUFB8^71^, which did not pass the significance threshold in our TurboID experiments but has a high-scoring phosphosite for both domains. Finally, we note that some potential binding sites were on proteins that were selectively observed in SHP2 mutant interactomes but not the SHP2^WT^ dataset. These include some members of respiratory complexes, as seen above, as well as Hsp40-family co-chaperones, which we discuss in the next section. Our interactomics data spanning multiple complexes of the electron transport chain expand the scope SHP2 interactors in the mitochondria and suggest plausible roles in this subcellular compartment.

### Destabilization of SHP2 protein structure might contribute to mitochondrial import

The mechanism by which some SHP2 mutants increasingly localize to the mitochondria is unclear. The Tom40 complex has been shown to mediate SHP2 translocation^69^. SHP2^WT^ and most mutants proximity-labeled Tom40 more than the TurboID-only control, but we did not observe mutant-specific enhancement of Tom40 engagement (**Figure 6A**). Tom20, a component of the Tom40 complex, binds mitochondrial localization sequences on cytoplasmic proteins, often at their N-termini^78^. SHP2 has a mitochondrial localization motif near its N-terminus (R4 to H8)^69^, and mutation of Arg4 and Arg5 to alanine residues diminishes mitochondrial localization (**Figure 5C**)^69^. Furthermore, in a recent Tom20 proximity-labeling study, SHP2 was identified as a proximal protein^79^. Notably, in our datasets, Tom20 is an enriched protein for several SHP2 mutants but not SHP2^WT^, as are other components of the Tom20/Tom40 complex (**Figure 6B and Supplementary Figure 8A**). We hypothesized that some SHP2 mutations might enhance exposure of the mitochondrial localization signal due to changes in protein conformation (**Figure 6C**). We analyzed our previously-reported molecular dynamics simulations of SHP2 in both the auto-inhibited state and an open state that is likely accessed by some disease mutants. These simulations showed an increase in solvent accessibility for R4 and F7 in the open conformation, but not for R5 (**Figure 6D**)^30^. Based on this analysis, it is plausible but not conclusive that SHP2 mutations alter Tom20 access to the mitochondrial localization sequence.

**Figure 6.**
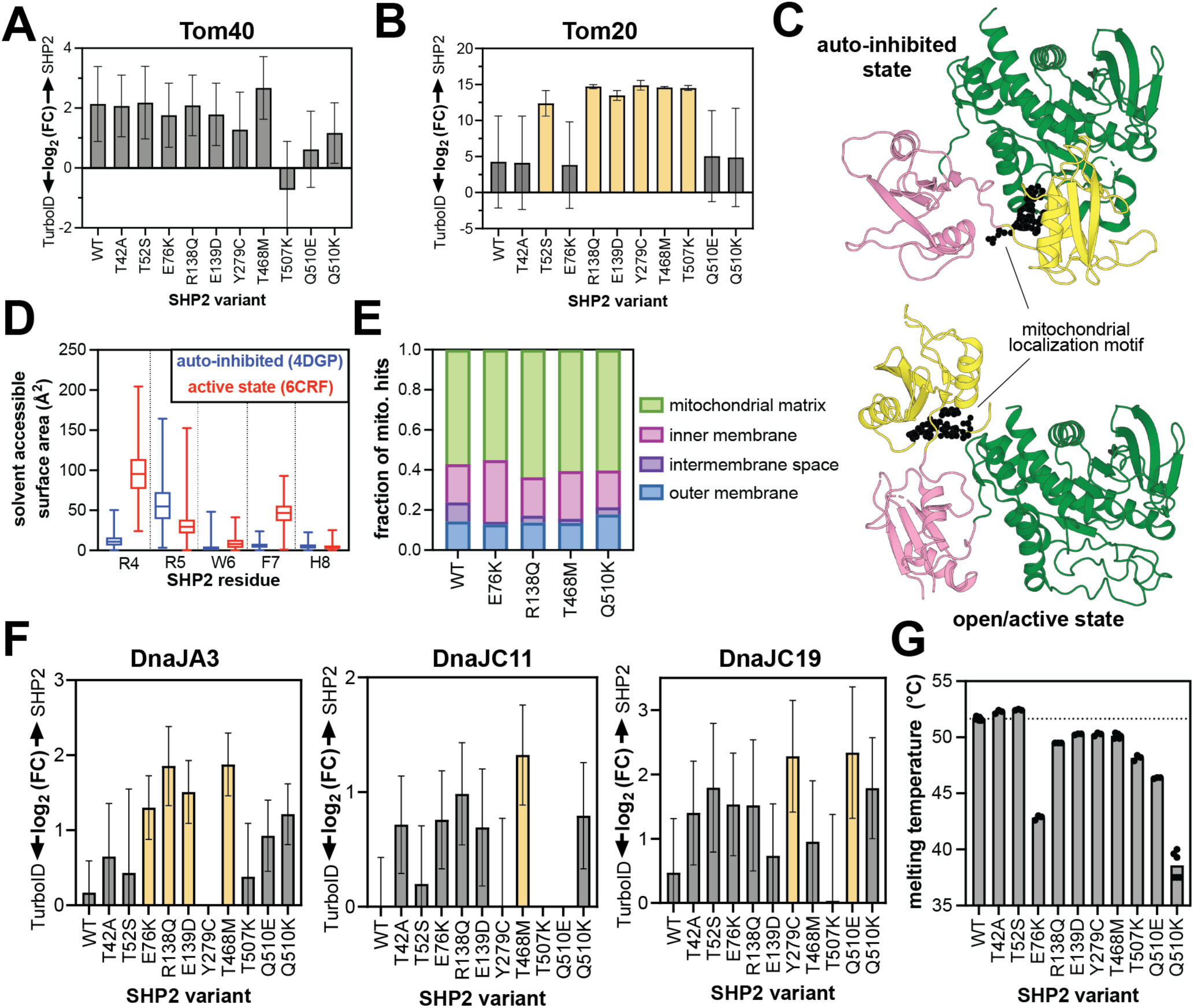
Protein interactions and structural features impacting SHP2 mitochondrial import. Proximity labeling of (**A**) Tom40 and (**B**) Tom20 outer mitochondrial import machinery. Variants in yellow are significantly enriched over the TurboID control (fold-change >2, p-value <0.05). (**C**) The SHP2 mitochondrial localization motif (R4 to H8) highlighted as black spheres on auto-inhibited (*left*, PDB code 4DGP) and open (*right*, PDB code 6CRF) structures. (**D**) Differences in solvent accessibility of the RRWFH motif in molecular dynamics simulations using auto-inhibited conformation (starting structure PDB code 4DGP) or open conformation (starting structure PDB code 6CRF). (**E**) Sub-mitochondrial localization of SHP2 variants inferred from proximity-labeling patterns. (**F**) Proximity-labeling signal for mitochondrial Hsp40/DnaJ proteins, DnaJA3, DnaJC11, and DnaJC19. Variants in yellow are significantly enriched over the TurboID control (fold-change >2, p-value <0.05). (**G**) Melting temperatures of SHP2 variants measured by differential scanning fluorimetry.

SHP2 has been observed both in the intermembrane space and the mitochondrial matrix, but most reported functions are in the matrix^65,69,71,77^. Our proteomics datasets show that SHP2^WT^ labels intermembrane space proteins more than the mutants that have increased overall mitochondrial signal, whereas those mutants mostly label matrix-associated proteins (**Figure 6E**). This suggests that SHP2^WT^ accumulates in the intermembrane space, or that some mutants experience more efficient transport through the inner mitochondrial membrane. Transport of soluble proteins through the inner membrane is typically mediated by the Tim23 complex. While components of this complex are present in our datasets, most do not show strong signal, other than Tim17B (**Supplementary Figure 8B**).

Mitochondrial transport and protein maturation in the mitochondrial matrix are highly dependent on chaperone proteins, most notably the Hsp70/40 system^80^. SHP2 is a known Hsp70 client^81^, however, neither cytoplasmic Hsp70 (HSPA1A/HSPA1B), nor its mitochondrial counterpart mortalin (HSPA9) were significantly enriched for any SHP2 variant in our datasets (**Supplementary Figure 8C**). However, multiple Hsp40/DnaJ proteins showed enhanced proximity-labeling signal for SHP2 mutants, most notably DnaJA3, DnaJC11, and DnaJC19, all of which play critical roles in the mitochondria (**Figure 6F**)^82^. DnaJC19 coordinates with Tim23 to facilitate protein translocation through the inner mitochondrial membrane^83^, DnaJA3 is critical for mitochondrial matrix proteostasis^84^, and DnaJC11 is involved in cristae formation and protein import^85^. It is noteworthy that, like SHP2, these proteins have been connected to significant developmental disorders and cardiomyopathies^86,87^. Our observations suggest that interactions with Hsp40 proteins could modulate the mitochondrial activity of SHP2 mutants.

SHP2 likely interacts with Hsp40/DnaJ proteins to facilitate its unfolding and refolding as it traverses the mitochondrial membranes, and these interactions presumably depend on the stability of SHP2. The auto-inhibited state of SHP2 has extensive interdomain interactions, likely making it more thermally stable than the SHP2 open state. Conversely, SHP2 mutants that adopt a more open conformation tend to have lower melting temperatures by differential scanning fluorimetry^7,30,88,89^. We previously showed that SHP2^R138Q^ and SHP2^T468M^ have lower melting temperatures than SHP2^WT^ and show here that SHP2^Q510K^ also has a decreased melting temperature (**Figure 6G**)^7,30^, suggesting that these mutants have a destabilized auto-inhibited state and/or decreased overall thermal stability. Collectively, our proximity-labeling data and stability measurements paint a plausible picture for mutation-dependent SHP2 translocation to the mitochondrial matrix, dictated by changes in protein stability and association with mitochondrial chaperones.

### Allosteric inhibition remodels the SHP2 interactome by stabilizing the auto-inhibited state

Our results illustrate how SHP2 conformation and stability can affect its interactome and localization. Several allosteric inhibitors of SHP2 are being pursued as cancer therapies, and many of these molecules bind between the C-SH2 and PTP domains to stabilize the auto-inhibited state (**Figure 7A**)^90,91^. One such inhibitor, TNO155 (batoprotafib), is currently in clinical trials^92^. Given its effect on SHP2 conformation and stability, we examined how TNO155 alters the SHP2 interactome. We pre-incubated SHP2^WT^-TurboID-transfected cells for 4 hours with 1 µM TNO155 prior to conducting proximity-labeling proteomics experiments. TNO155 treatment dramatically reduced the number of interactors for SHP2^WT^ in unstimulated and EGF-stimulated cells (**Figure 7B,C**, **Supplementary Table 2**). These results are consistent with auto-inhibited SHP2 being less capable of engaging in protein-protein interactions.

**Figure 7.**
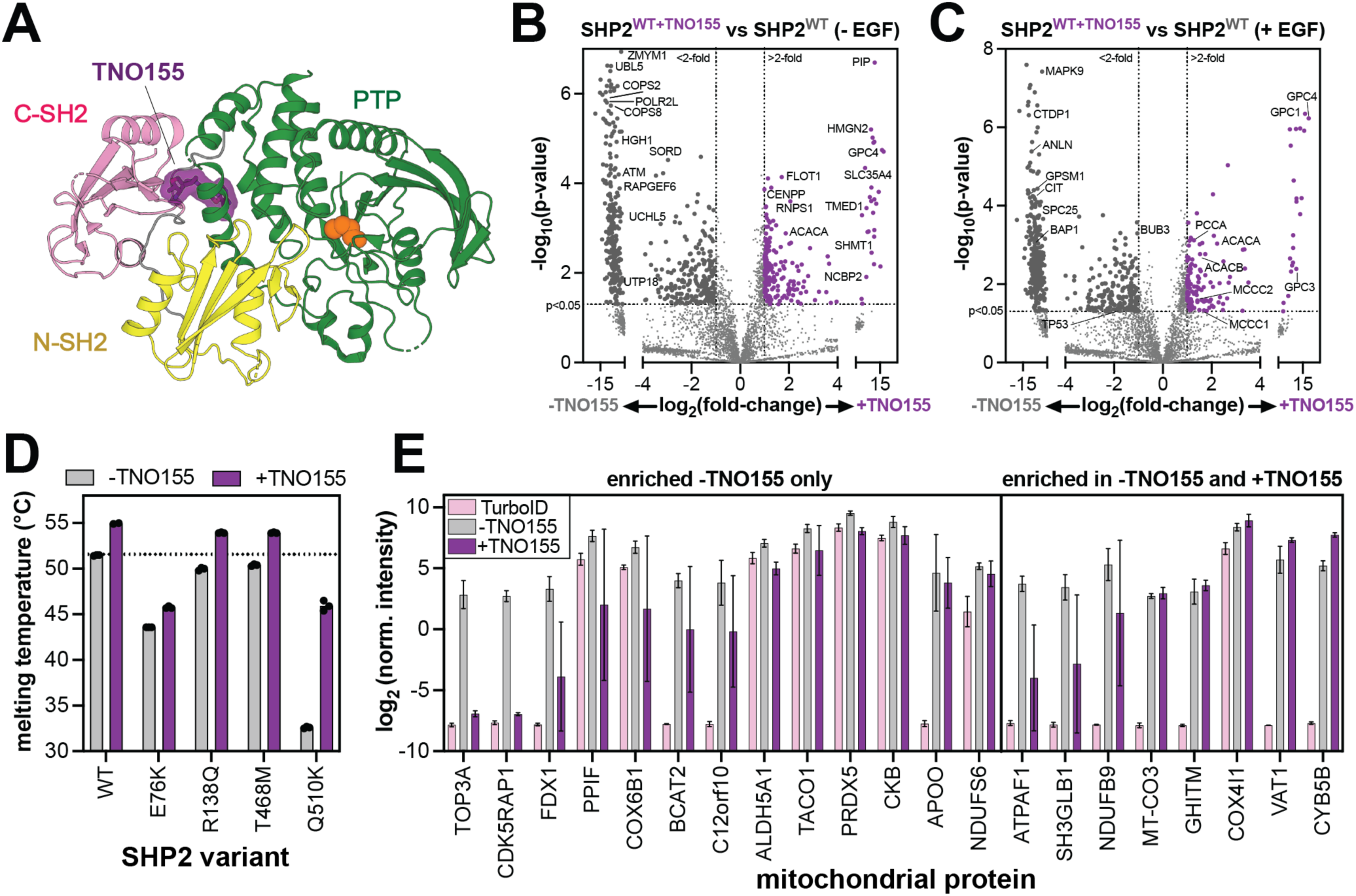
Alteration of SHP2 interactions by the allosteric inhibitor TNO155. (**A**) Structure of SHP2 with allosteric inhibitor TNO155 docked in between the C-SH2 domain and the PTP domain (PDB code 7JVM). (**B**) Volcano plot highlighting enriched proteins in SHP2^WT^, compared to SHP2^WT+TNO155^ in the absence of EGF stimulation. 210 proteins were significantly enriched in inhibitor-treated samples over untreated samples, and 480 proteins were significantly enriched in untreated samples over inhibitor-treated ones. (**C**) Volcano plot highlighting enriched proteins in SHP2^WT^, compared to SHP2^WT+TNO155^ in the presence of EGF stimulation (**D**) Melting temperatures of SHP2 variants measured by differential scanning fluorimetry in absence and presence of 10 μM TNO155. (**E**) Normalized intensities for 21 mitochondrial proteins that are significantly enriched in SHP2^WT^ versus TurboID-only. Intensities are shown for the TurboID-only control (pink), SHP2^WT^ without TNO155 (gray), and SHP2^WT^ with TNO155 (purple). The 13 proteins on the left are not significantly enriched in the +TNO155 sample over the TurboID-only control. The 8 proteins on the right are significantly enriched in the +TNO155 sample over the TurboID control.

Given that TNO155 enforces the auto-inhibited state, we hypothesized that it would enhance the thermal stability of SHP2, which in turn could impact SHP2 mitochondrial import. We first measured the melting temperatures of SHP2^WT^ and mutants with enhanced mitochondrial localization in the absence and presence of TNO155 (**Figure 7D**). Each SHP2 variant showed an increase in melting temperature upon addition of TNO155, albeit to varying extents. This variability likely reflects the extent to which SHP2 adopts an open conformation that lacks the inhibitor binding site. Indeed, SHP2^E76K^, which primarily inhabits the open conformation showed the smallest increase of ∼2 °C^61,93^. By contrast, SHP2^Q510K^ showed a dramatic increase of 13 °C, but the molecular basis for this large effect is not immediately obvious. These results show that TNO155 binding can enhance the stability of SHP2.

Next, we examined the mitochondrial proteins enriched in SHP2^WT^-TurboID experiments with and without TNO155. As previously noted, we identified 78 proteins enriched in SHP2^WT^ in unstimulated cells, 21 of which were mitochondrial proteins (**Figure 5A**). Upon TNO155 treatment, 13 of these mitochondrial proteins were no longer significantly enriched over the TurboID-only control (**Figure 7E**, *left*), supporting the notion that SHP2 stability is a driving factor in its subcellular localization. Remarkably, a few mitochondrial SHP2^WT^ interactors were still enriched upon TNO155 treatment (**Figure 7E**, *right*). These proteins could be interactors that engage SHP2 at inhibitor-insensitive interfaces, or they might be co-localized proteins but not direct interactors. . Alternatively, these proteins might bind tightly to SHP2 in the open state, rendering it insensitive to TNO155, as has been shown for other SHP2 inhibitors and interactors^62^. Notably, both NDUFB9 and ATPAF1 have predicted high-affinity N- or C-SH2 binding sites (**Figure 5F**, **Supplementary Table 4**). These different mechanisms warrant further investigation, as they could have implications for inhibitor efficacy. Collectively, these results show that the allosteric inhibitor TNO155 alters both SHP2 interactions and its localization by changing its overall conformation and stability. These results also highlight the exquisite sensitivity and broad applicabiliy of TurboID proximity labeling to examine state-dependent changes in SHP2 function.

## Discussion

Disease-driving mutations in *PTPN11*/SHP2 are almost exclusively missense mutations, many of which disrupt auto-inhibition and hyperactivate SHP2^10^. Characterization of these mutations has broadened our understanding of the protein-intrinsic regulation of SHP2 and shed light on how dysregulation of SHP2 can cause disease. While SHP2 functions in many signaling pathways, its role in disease has largely been linked to dysregulation of the Ras/MAPK pathway^13,14,94–96^. Recent studies have started to reveal alternate mechanisms of SHP2 dysregulation and pathogenicity. For example, some catalytically inactive NSML mutants drive proliferative signaling through increased adoption of an open conformation, which enhances SH2-phosphoprotein interactions^31,97^. Other mutations alter substrate- or ligand-binding preferences and rewire signaling interactions^7,34^. This range of mutational effects motivated us to systematically map the cellular interactomes of diverse SHP2 mutants to gain insights into resulting aberrant cell signaling.

In this study, we show that TurboID proximity labeling allows for in-depth comparison of mutation- and drug-induced changes in SHP2 protein-protein interactions and localization. Our results show that individual SHP2 mutants have overlapping but partly distinct interactomes that reveal both divergent and convergent modes of dysregulation. For example, the melanoma mutant SHP2^R138Q^ has a dysfunctional C-SH2 domain, and interactions with this mutant are less dependent on tyrosine-phosphorylated proteins than SHP2^WT^. The Noonan Syndrome mutant SHP2^T42A^ enhances intrinsic phosphoprotein binding affinity and alters binding specificity, increasing its interaction with the cell surface receptor MPZL1. SHP2^T42A^ biochemically phenocopies NSML mutants SHP2^T468M^ and SHP2^Q510E^, which adopt an open conformation that also stabilizes their interaction with MPZL1. This observation highlights how mechanistically distinct mutations can yield similar cellular and clinical phenotypes.

A key aspect of our study is the juxtaposition of SHP2 proximity-labeling data with predictions of SH2 binding sites, revealing a number of tyrosine phosphorylation sites that could recruit and modulate SHP2 function. The PTP domain of SHP2 also interacts with phosphosites, to dephosphorylate them. From our proximity-labeling datasets alone, we cannot differentiate binders and co-localized proteins from direct substrates, and the SHP2 PTP domain likely does not have strong sequence preferences that could be used to predict substrates^43^. Recently, substrate-trapping mutations in PTP1B, another phosphatase, were successfully combined with proximity labeling to selectively enhance signal for substrates over other proximal proteins^98^. However, certain SHP2 substrate-trapping mutations, most notably C459S, cause dramatic structural rearrangements that will have collateral effects on SHP2 localization and the measured interactome^61,99^. Thus, careful assessment of the structural effects of substrate-trapping mutants will be required.

A notable feature of our proximity-labeling datasets is that we observed a strong mitochondrial signal for SHP2^WT^ and several mutants. The role of SHP2 in mitochondrial biology is not well-understood. Hyperactive SHP2 variants can dysregulate mitochondrial function, causing oxidative stress in cells^33,68,77^. SHP2^WT^ regulates the inflammasome through interaction with ANT1, and SHP2^E76K^ can enhance mitochondrial metabolism through excessive dephosphorylation of STAT3^69,71^. Here, we show that several SHP2 variants co-localize with components of the electron transport chain. Furthermore, we show that SHP2^R138Q^, SHP2^T468M^ and SHP2^Q510K^, and SHP2^E76K^ have increased mitochondrial localization relative to SHP2^WT^, and some of these mutants can alter mitochondrial respiration. Our data suggest that some mutants have enhanced interactions with mitochondrial import machinery or associated chaperones, due to a change in protein conformation or stability. This is further supported by our analysis of the SHP2^WT^ interactome in presence of the allosteric inhibitor TNO155, which stabilizes SHP2 in the auto-inhibited state and reduces mitochondrial proximity-labeling signal for SHP2^WT^. However, we note that some SHP2 mutants with decreased thermal stability do not appear to have increased mitochondrial localization, suggesting that there is nuance to the molecular mechanism of SHP2 mitochondrial import. These observations motivate further studies probing SHP2 mislocalization and mitochondrial dysregulation. More broadly, our work lays the foundation for new avenues of investigation into SHP2 signaling and pathogenicity. Ultimately, we envision that the experimental framework laid out in this study will be useful for the unbiased dissection of mutational effects in other disease-relevant proteins.

## Acknowledgements

We would like to thank the members of the Shah and Jovanovic labs for their scientific insights and helpful discussions. This research was funded by NIH/NIGMS grant R35GM138014 to NHS. MJ was supported by NIH/NIGMS grant R35GM152258. JTG was funded by NIH/NCI grant 1DP2CA281605. LCT is supported by an NSF Graduate Research Fellowship (award # 2036197).

## Author contributions

AEV conceived, designed, performed, analyzed, and interpreted the experiments; performed statistical analysis; and wrote the manuscript. LCT designed experiments and advised on interpretation and analysis of the data. ZJ and RV performed experiments. AI performed, analyzed, and interpreted the experiments. KK analyzed the data and interpreted the experiments. JTG contributed key resources. MJ designed and interpreted the experiments and edited the manuscript. NHS designed, analyzed, and interpreted the experiments, and wrote the manuscript.

## Competing interests

The authors declare no competing interests.

## Data and materials availability

All of the processed data, including log_2_ intensities from the AP-MS, TurboID-MS and corresponding total proteomes, and the enrichment scores from the high-throughput SH2 specificity screens are provided as supplementary table files. The microscopy images and deep-sequencing data from specificity screens are available as a Dryad repository (DOI: 10.5061/dryad.wstqjq2xv). Raw mass spectrometry data will be made available via ProteomeXchange. Plasmids and DNA libraries generated in this study will be made freely available upon request. There are no restrictions to the availability of data and reagents generated in this study.

## Notes

https://doi.org/10.5061/dryad.wstqjq2xv

https://www.ebi.ac.uk/pride/archive/projects/PXD067419

https://www.ebi.ac.uk/pride/archive/projects/PXD067905

### Competing Interest Statement

The authors have declared no competing interest.

### Summary of Updates

1. We updated the supplementary table files to include a clearer legend on the first tab of every spreadsheet. 2. We updated funding information, which was incomplete in the prior versions. 3. We identified an error in the sample labeling in Supplementary Table 3. This has now been fixed, and associated Supplementary Figure 2G,H have been updated as well. 4. There was an error in both the cropping and resulting interpretation of the blot in Supplementary Figure 2F. The figure panel has been updated, and a sentence in the main text on page 5 has been revised, accordingly.

