## Supplementary Information and Figures for "Proximity-labeling proteomics reveals remodeled interactomes and altered localization of pathogenic SHP2 variants"

**Table of contents:**

Supp. Fig. 1. Affinity-purification mass spectrometry with SHP2$^{WT}$ and SHP2$^{T42A}$ pg 2
Supp. Fig. 2. Quality control and validation of the SHP2-TurboID system pg 3
Supp. Fig. 3. STRING interaction networks for SHP2$^{WT}$-TurboID hits pg 4
Supp. Fig. 4. Mutation- and EGF-dependent changes in SHP2-TurboID proximity labeling pg 5
Supp. Fig. 5. STRING interaction networks for core SHP2 interactomes pg 6
Supp. Fig. 6. Mitochondrial localization of SHP2 pg 7
Supp. Fig. 7. SHP2 proximity-labeling distributions in subcellular compartments pg 8
Supp. Fig. 8. Mutant-specific labeling of mitochondrial import proteins and chaperones pg 9
Materials and methods pg 10
Supplementary references pg 18

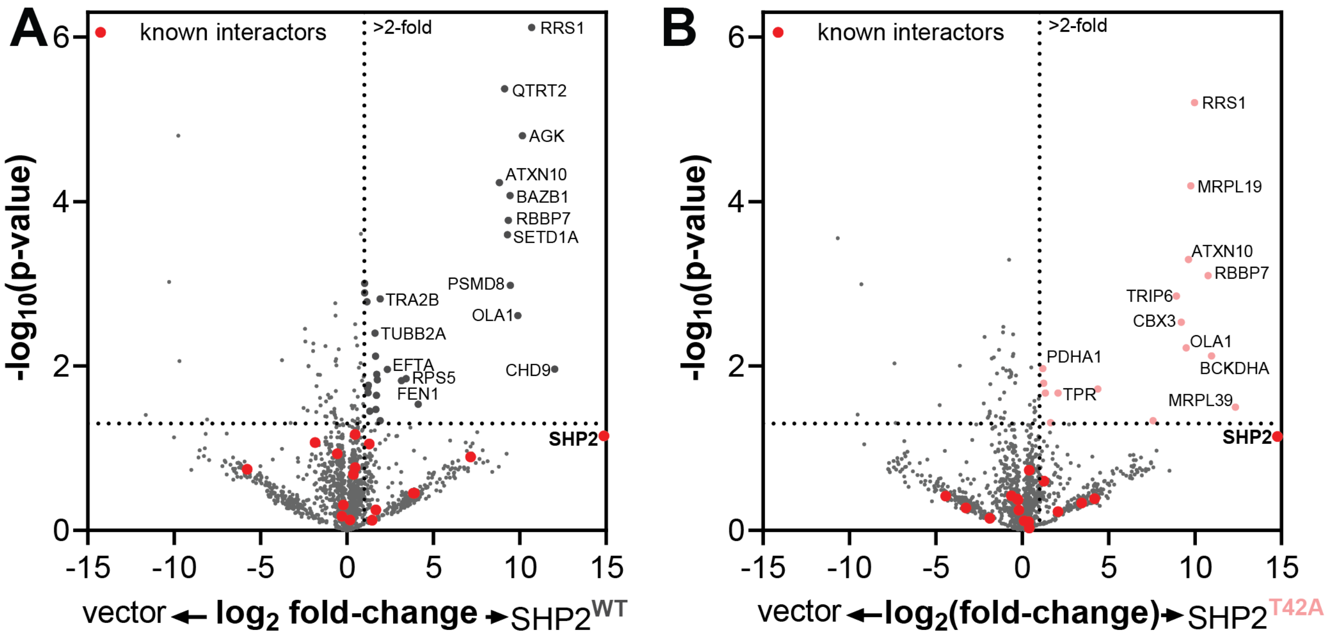


**Supplementary Figure 1. Affinity-purification mass spectrometry with SHP2$^{WT}$ and SHP2$^{T42A}$.** (**A**) Volcano plot showing proteins enriched in SHP2$^{WT}$ over a vector control using affinity purification mass spectrometry. No known positives were identified (n = 3). (**B**) Same as (**A**), but for SHP2$^{T42A}$ (n = 3).

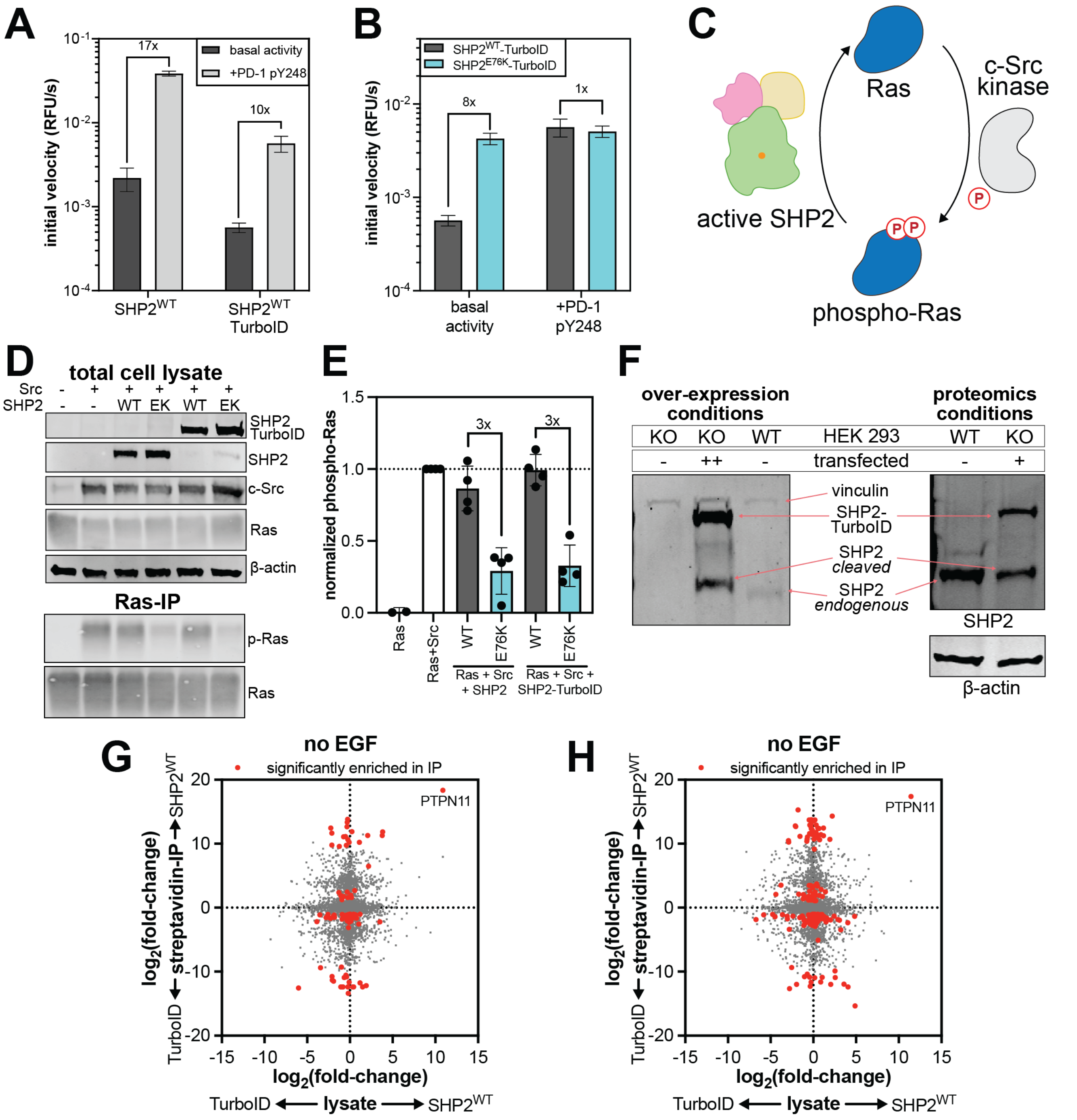


**Supplementary Figure 2. Quality control and validation of the SHP2-TurboID system.** (**A**) *In vitro* activity measurements showing that SHP2-TurboID can be activated by a phosphopeptide (PD-1 pY248) (n = 3). (**B**) *In vitro* activity measurements showing that E76K hyperactivation is preserved in a TurboID-fusion context, and only $SHP2^{WT}$-TurboID, but not $SHP2^{E76K}$-TurboID can be further activated by PD-1 pY248 (n = 3). (**C**) Schematic of the Ras dephosphorylation assay in HEK 293 cells. (**D**) Representative western blots showing dephosphorylation N-Ras by SHP2-TurboID proteins. WT = $SHP2^{WT}$ EK = $SHP2^{E76K}$. (**E**) Quantification of Ras dephosphorylation assays (n=4). (**F**) The left blot compares SHP2 levels in WT and $SHP2^{KO}$ HEK 293 cells, and shows over-expression of $SHP2^{WT}$-TurboID in $SHP2^{KO}$ cells, highlighting partial cleavage of SHP2-TurboID. The right blot compares endogenous SHP2 levels in WT HEK 293 cells with $SHP2^{WT}$-TurboID levels in $SHP2^{KO}$ HEK 293 cells, transfected under the same conditions used for proteomics experiments. (**G**) Fold-change between $SHP2^{WT}$-TurboID and TurboID-only for total lysates, measuring protein abundance (x-axis), and streptavidin-IP, measuring proximity labeling (y-axis), in unstimulated cells. (**H**) Same as (**G**), but for cells stimulated with 100 ng/mL EGF (n = 3 for total lysate and TurboID datasets).

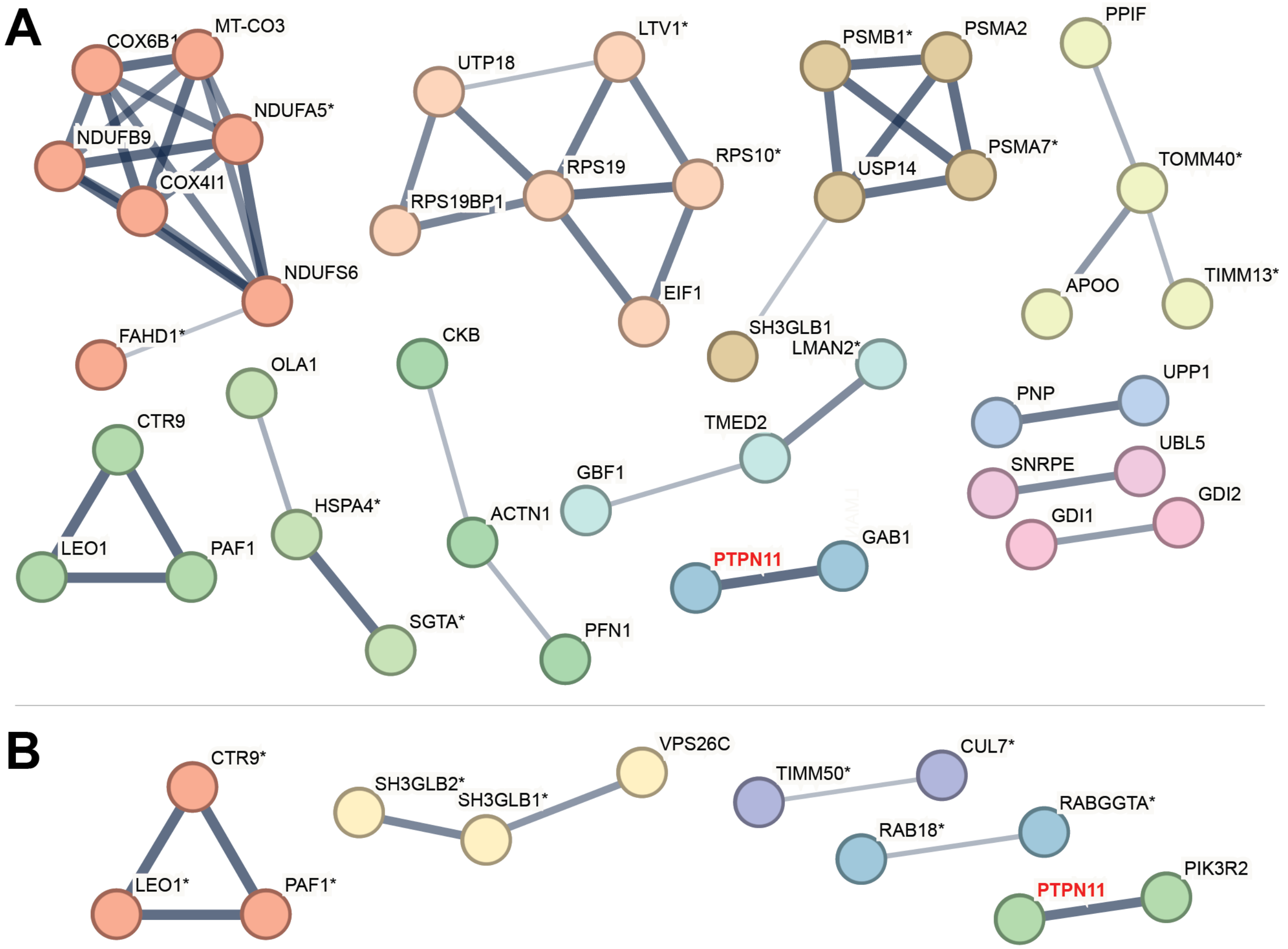


**Supplementary Figure 3. STRING interaction networks for SHP2$^{WT}$-TurboID hits.** (**A**) STRING interaction network for proteins enriched by SHP2$^{WT}$-TurboID over the TurboID control by at least 2-fold with a p-value <0.1, in the absence of EGF stimulation. (**B**) same as in (**A**) but with EGF stimulation. Proteins with a p-value between 0.05 and 0.1 are marked with an asterisk. All other proteins have a p-value <0.05. For both panels, solid lines between proteins indicate a known physical interaction, as documented in the STRING database. Only proteins that have a physical interaction with at least one other protein in our interactomes and have an edge confidence score of at least 0.4 are shown. Edge thickness represents edge confidence: thin = 0.4, medium = 0.7, thick = 0.9. Clusters were identified by Markov Clustering (MCL).

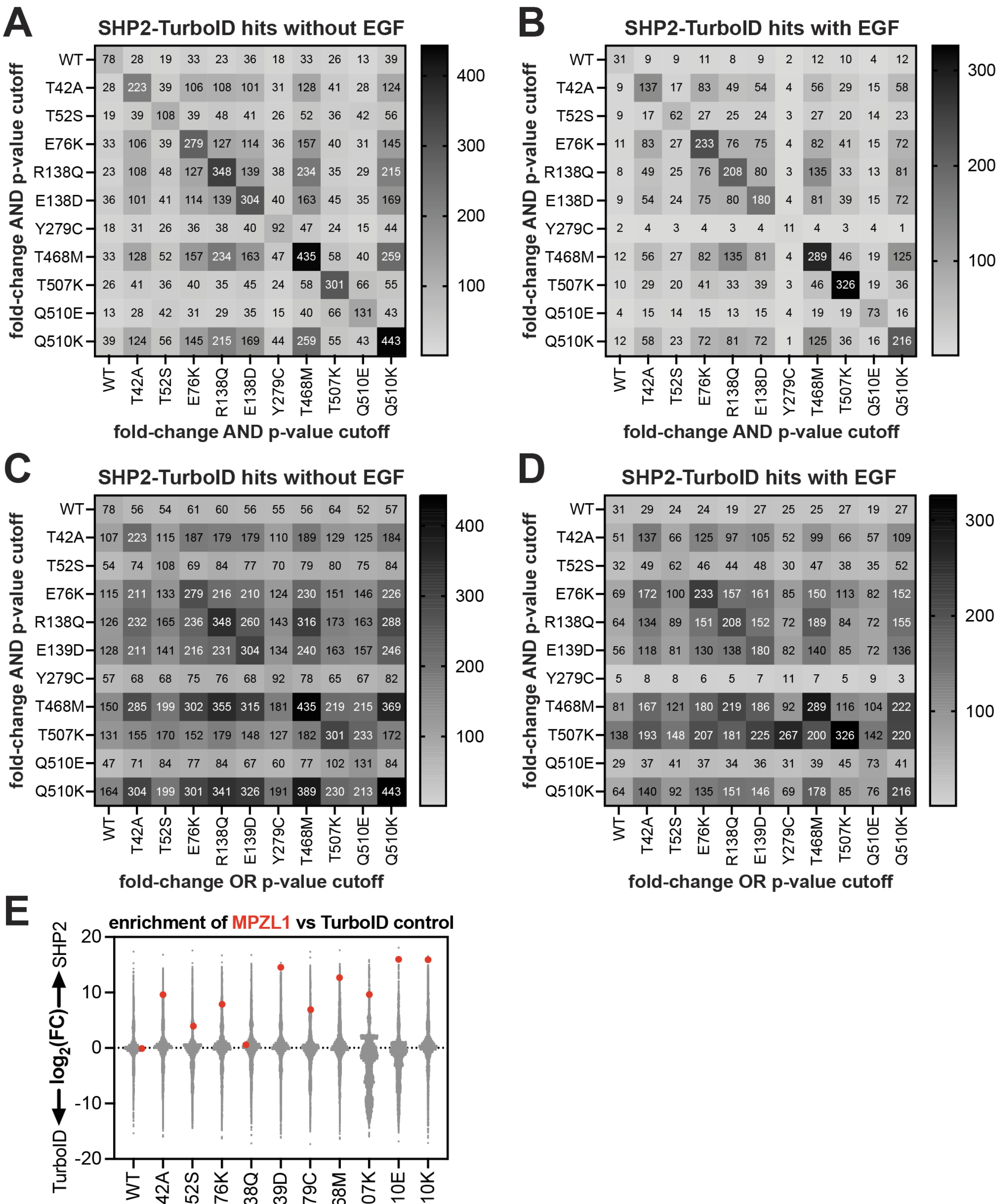


**Supplementary Figure 4. Mutation- and EGF-dependent changes in SHP2-TurboID proximity labeling.** (**A**) Stringent pairwise overlap in hits for all SHP2 variants relative to the TurboID control in unstimulated samples. Hits are defined as proteins with >2-fold enrichment **and** a p-value <0.05, and overlap indicates the number of proteins that meet these criteria for both SHP2 variants in the comparison. (**B**) Same as in (**A**) but with EGF stimulation. (**C**) Relaxed pairwise overlap in hits for all SHP2 variants relative to TurboID control in unstimulated samples. For SHP2 variants on the y-axis, a hit was defined as proteins with both >2-fold enrichment **and** a p-value <0.05. For SHP2 variants on the x-axis, a hit was defined as proteins with either >2-fold enrichment **or** a p-value <0.05. (**D**) Same as (**C**), but with EGF stimulation. Note that in panels (**A**)-(**D**) the smaller numbers for SHP2$^{Y279C}$ reflect the use of only 2 replicates for this mutant, due to an instrument error and sample loss. (**E**) Distribution of protein enrichments for SHP2 variants versus the TurboID control, highlighting MPZL1 enrichment by some mutants.

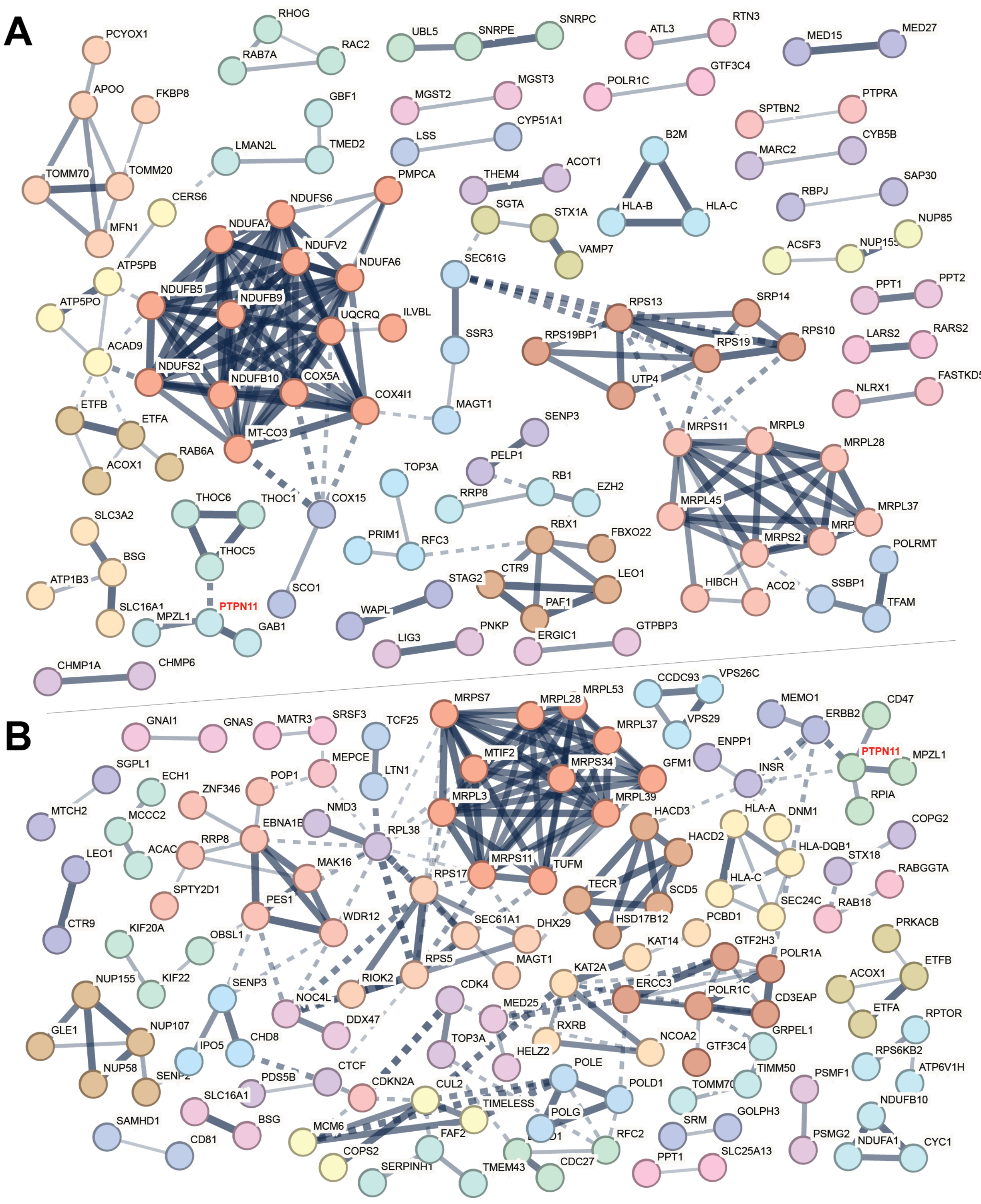


**Supplementary Figure 5. STRING interaction networks for core SHP2 interactomes.** (**A**) STRING interaction network for proteins significantly enriched by 4 or more SHP2 variants in the absence of EGF stimulation. (**B**) STRING interaction network for proteins significantly enriched by 3 or more SHP2 variants in the absence of EGF stimulation. For both panels, solid lines between proteins indicate a known physical interaction between proteins within a cluster, as documented in the STRING database. Dashed lines also indicate a known physical interaction, but for proteins between two clusters. Only proteins that have a physical interaction with at least one other protein in our interactomes and have an edge confidence score of at least 0.4 are shown. Edge thickness represents edge confidence: thin = 0.4, medium = 0.7, thick = 0.9. Clusters were identified by Markov Clustering (MCL).

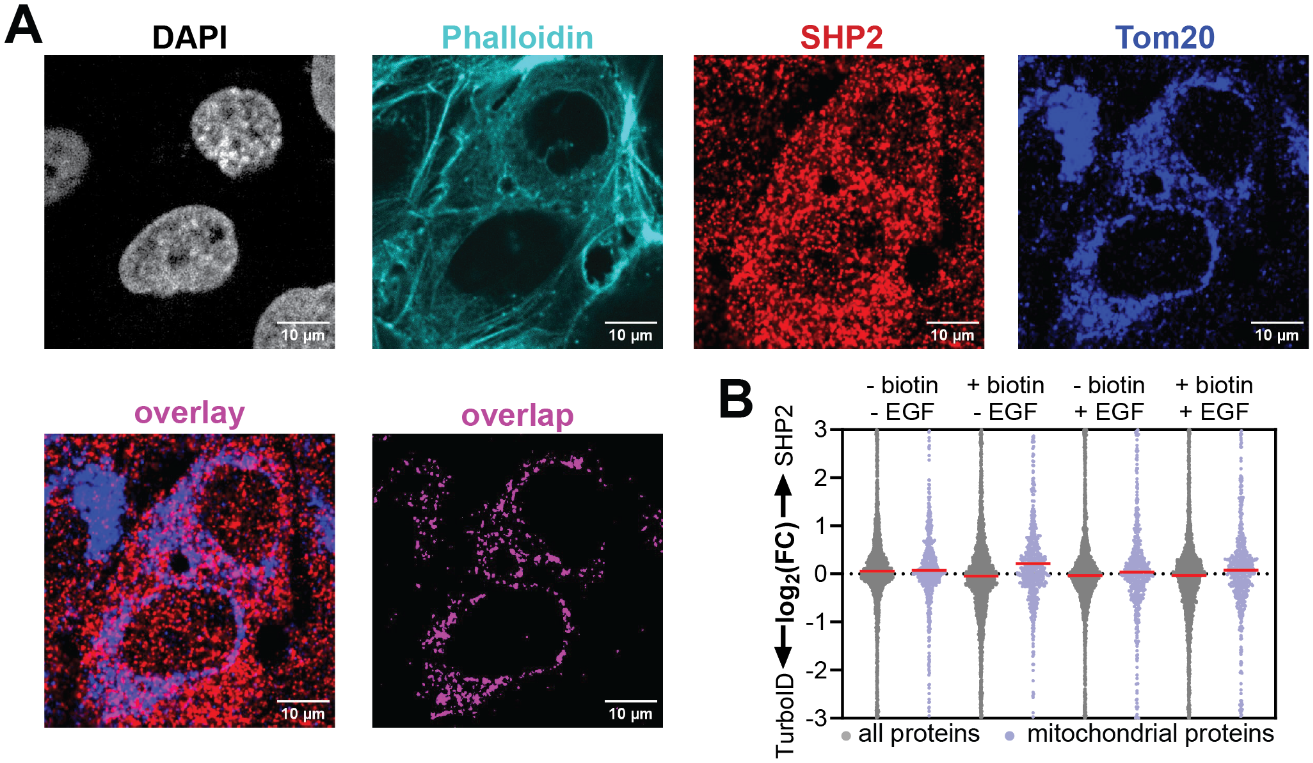


**Supplementary Figure 6. Mitochondrial localization of SHP2.** (**A**) Analysis of endogenous SHP2$^{WT}$ localization in U2-OS cells by confocal fluorescence microscopy showing SHP2 mitochondrial localization. (**B**) Enrichment of all proteins versus annotated mitochondrial proteins in SHP2$^{WT}$-TurboID samples compared with TurboID alone, with and without exogenously supplied biotin.

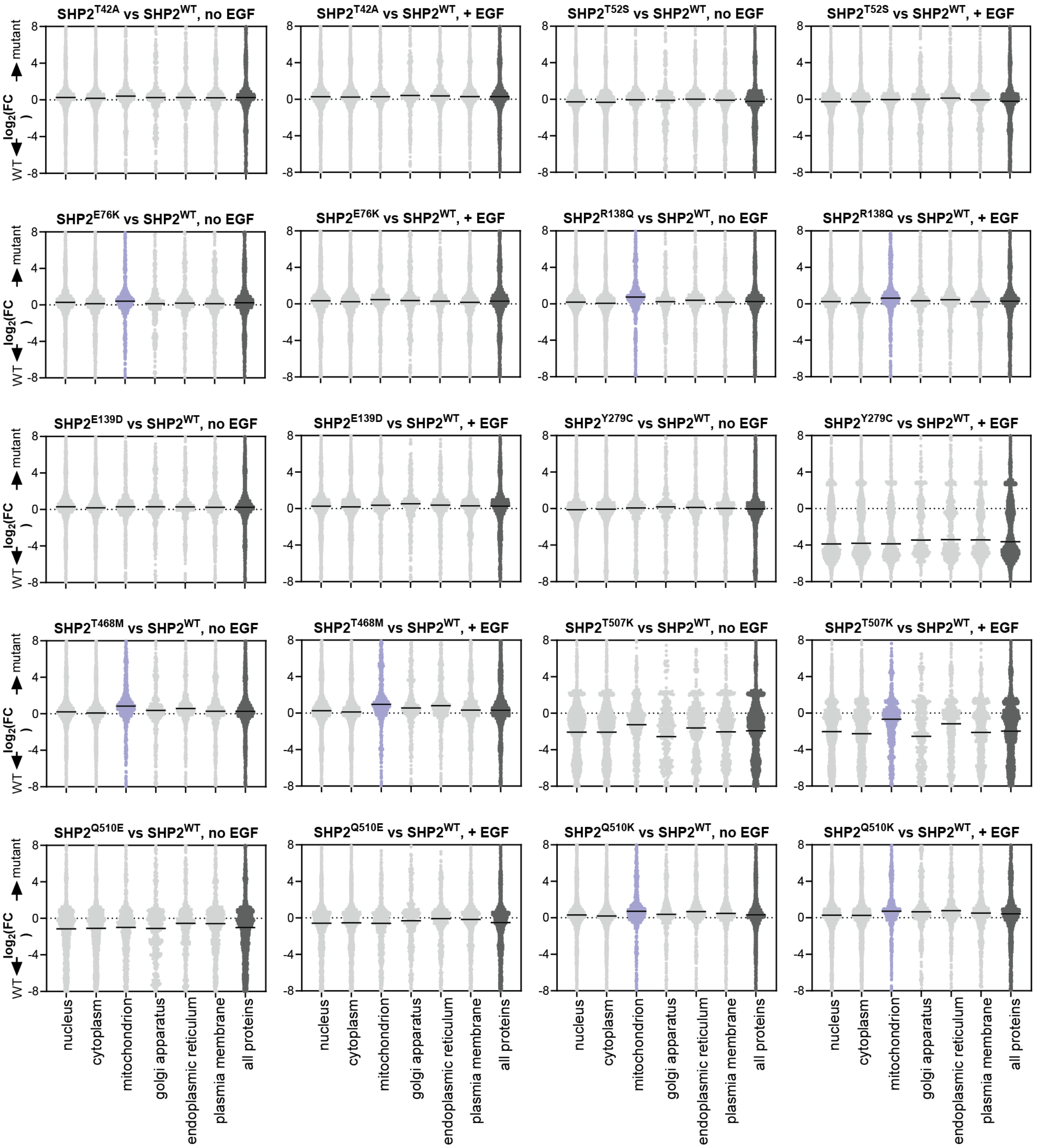


**Supplementary Figure 7. SHP2 proximity-labeling distributions in subcellular compartments.** Each graph shows a series of distributions made up of proteins found in different subcellular compartments. The distributions show the enrichment values for those proteins in SHP2-TurboID samples relative to the TurboID control. Subcellular distributions with a distinctive increase relative to the “all proteins” distribution (dark gray) are highlighted in color.

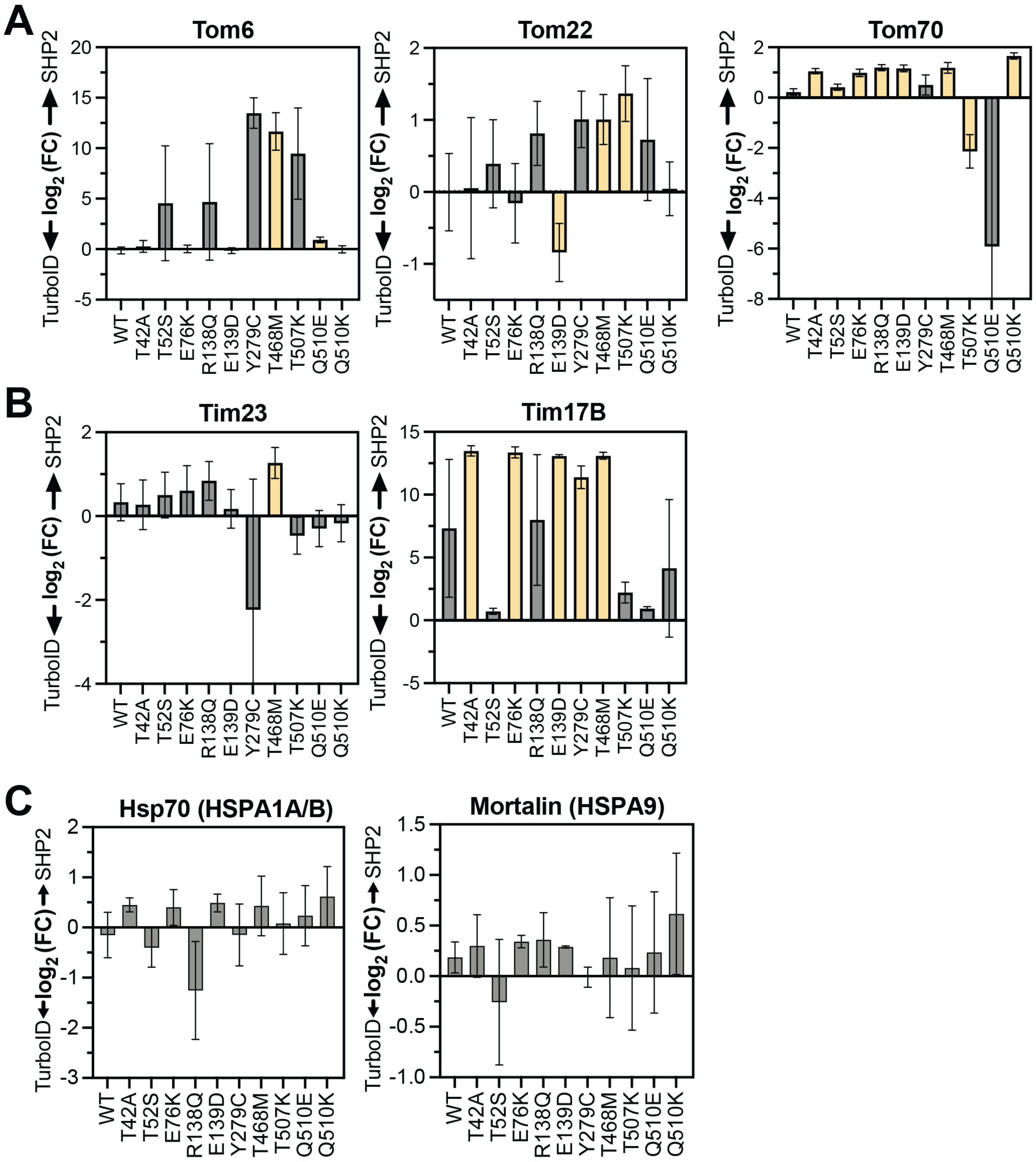


**Supplementary Figure 8. Mutant-specific labeling of mitochondrial import proteins and chaperones.** (**A**) Enrichment or depletion of select Tom proteins in our dataset for all SHP2 variants relative to the TurboID-only control. Bars in yellow indicate a significant difference from the TurboID-only control (fold-change >2, p-value <0.05). (**B**) Same as (**A**), but for two components of the Tim23 complex in our dataset. (**C**) Same as (**A**) but for HSPA1A/HSPA1B (cytosolic Hsp70) and HSPA9/mortalin (mitochondrial Hsp70).

### Materials and methods

#### Cell culture

All cell lines were cultured in a 37 °C tissue culture incubator with 5% $CO_2$. Cells were discarded by passage 25 and tested for mycoplasma every 3 months. HEK 293 cells were grown in Dulbecco's Modified Eagle Medium (DMEM) with 10% Fetal Bovine Serum (FBS) and 1% penicillin/streptomycin. HEK 293 SHP2 knock-out and U-2 OS cells, reported previously[1], were grown in DMEM with 10% FBS. Human epidermal growth factor was purchased in lyophilized form (#E9644, Sigma) and reconstituted in 10 mM acetic acid.

#### Myc-tag immunoprecipitation for proteomics

Replicates consist of separate transfections and downstream processing. 10 x $10^6$ HEK 293 SHP2 KO cells were seeded in a 15 cm plate. The next day, cells were transfected overnight with 25 µg DNA (Myc-tagged SHP2-WT, SHP2-T42A or empty pEF vector) in 2.5 mL empty DMEM using 75 µg of polyethylenimine (PEI). The next morning, the transfection medium was replaced by pre-warmed empty DMEM. 36 hours after transfection, cells were harvested by scraping and washed twice in 1 mL phosphate buffered saline (PBS) pre-warmed to 37 °C. Half of the samples were resuspended in 1 mL PBS with 100 ng/mL epidermal growth factor (EGF), the other half was resuspended in 1 mL PBS. Samples were placed in a 37 °C heat block for 10 minutes and then placed on ice to stop the reaction. Cells were spun down in a refrigerated tabletop centrifuge. Cells were lysed in non-denaturing lysis buffer (20 mM Tris HCl pH 8.0, 137 mM NaCl, 10% glycerol, 1% Nonidet P-40 (NP-40), 2 mM EDTA, with protease inhibitors and phosphatase inhibitors added fresh) for 25 minutes at 4 °C while rotating, then spun down in a refrigerated tabletop centrifuge at 17,000 g for 15 minutes. Protein concentration was determined using a bicinchoninic acid (BCA) assay. 1500 µg of protein was used in an immuno-precipitation using 75 µL of magnetic anti-Myc beads. Samples were left overnight at 4 °C while rotating. The next day, samples were prepared for proteomics.

#### Preparation of TurboID samples

Replicates consist of separate transfections and downstream processing. 10 x $10^6$ HEK 293 $SHP2^{KO}$ cells were seeded in a 15 cm plate. The next day, cells were transfected overnight with 25 µg SHP2-TurboID DNA in 2.5 mL empty DMEM using 75 µg of PEI. The next morning, the transfection medium was replaced by pre-warmed empty DMEM with 2.5% FBS. 36 hours after transfection, media was aspirated and replaced by one of the following: empty DMEM (unstimulated control), empty DMEM with 100 ng/mL EGF (stimulated control), 100 µM biotin in DMEM (unstimulated sample), or 100 µM biotin and 100 ng/mL EGF in DMEM (stimulated sample). Cells were placed at 37 °C for 10 minutes and then promptly placed on ice. Cells were harvested in 10 mL of ice-cold PBS by scraping and washed twice in 5 mL cold PBS. Cells were lysed in non-denaturing lysis buffer (20 mM Tris HCl pH 8.0, 137 mM NaCl, 10% glycerol, 1% Nonidet P-40 (NP-40), 2 mM EDTA, with protease inhibitors and phosphatase inhibitors added fresh) for 25 minutes while rotating at 4 °C, then spun down in a refrigerated tabletop centrifuge at 17,000 g for 15 minutes. Protein concentration was determined using a bicinchoninic acid (BCA) assay. 1500 µg of protein was used in an immuno-precipitation using 75 µL of magnetic anti-Streptavidin beads. Samples were left overnight at 4 °C while rotating. The next day, samples were prepared for proteomics.

#### Preparation of proteomic IP-samples

Samples were prepared according to previously a reported protocol[2]. Beads were washed twice in 1 mL 50 mM Tris-HCl (pH 8.0), then twice more in 200 µL 2M urea in 50 mM Tris (pH 8.0). Beads were resuspended in80 µL of 2M urea in 50 mM Tris-HCl (pH 8.0), containing 1 mM DTT and 0.4 µg trypsin at room temperature while shaking moderately. After 1 hour, the supernatant was transferred to a new tube. Beads were washed twice with 60 µL of 2M urea in 50 mM Tris (pH 8.0). The washes were combined with the on-bead digest 80 µL supernatant to a total volume of 200 µL. Dithiothreitol (DTT) was added to a final concentration of 4 mM, and incubated at room temperature while shaking moderately. After 30 minutes, iodoacetamide (IAA) was added to a final concentration of 10 mM and incubated at room temperature in the dark while shaking moderately. After 45 minutes, an additional 0.5 µg of trypsin was added to each sample. Digestion proceeded overnight at room temperature while shaking. After overnight digestion, samples were acidified using formic acid (FA) to ~1% (vol/vol). Samples were desalted using C18 stage tips. Briefly, tips were conditioned with 100 µL of 100% MeOH, 100 µL of 0.2% FA/60% Acetonitrile (ACN), and twice with 100 µL of 0.2% FA. Acidified peptides were loaded onto the stagetips and washed twice with 100 µL of 0.2% FA. Peptides were eluted with 50 µL of 0.2% FA/60% ACN, and dried in a vacuum centrifuge at room temperature. Peptides were stored in the -80°C until injection.

### Preparation of lysate proteomic samples

50 µg of protein was used as input for the lysate samples and brought up to a total volume of 200 µL in 50 mM Tris-HCl (pH 8.0). Dithiothreitol (DTT) was added to a final concentration of 5 mM. Samples were incubated for 45 minutes at room temperature while shaking moderately. Iodoacetamide (IAA) was added to a final concentration of 10 mM and shaken for another 45 minutes in the dark at room temperature. 10 µL of magnetic SP3 beads at a concentration of 50 mg/mL were added to the samples. 250 µL of EtOH was added and samples were placed on a magnetic rack. Supernatant was aspirated and beads were washed three times for two minutes with 1mL 80% EtOH. Samples were reconstituted in 200 µL fresh 100 mM ammonium bicarbonate (ABC), pH 7.7. Trypsin was added in a 1:50 ratio and samples were digested overnight while lightly shaking. In the morning, the supernatant was transferred to a new tube and spun down at 17,000 g for 5 minutes. 175 µL of supernatant was transferred to another new tube and 1% FA (v/v) was added. Samples were dried down in a vacuum centrifuge at room temperature and dried peptides were stored in -80°C until injection.

### LC-MS/MS analysis on a Q-Exactive HF for immunoprecipitated samples

Samples were resuspended in 13 µL 3% ACN / 0.2% FA. 6 µL was injected and analyzed on a Waters M-Class UPLC using a 25cm Ionopticks Aurora column coupled to a benchtop Thermo Fisher Scientific Orbitrap Q Exactive HF mass spectrometer. Peptides were separated at a flow rate of 400 nL/min with a 100 min gradient, including sample loading and column equilibration times. Data was acquired in data dependent mode using Xcalibur 4.1 software. MS1 Spectra were measured with a resolution of 120,000, an AGC target of 3e6 and a mass range from 300 to 1800 m/z. Up to 12 MS2 spectra per duty cycle were triggered at a resolution of 60,000, an AGC target of 1e5, an isolation window of 0.8 m/z, a normalized collision energy of 28, a scan range of 200 to 2000 m/z, and a fixed first mass of 110 m/z.

### Quantification and statistical analyses of immunoprecipitated samples

All raw data were analyzed with SpectroMine software version 4.2.230428 using a UniProt database (Homo sapiens, UP000005640). Carbamidomethylation on cysteines was set as a fixed modification. Oxidation of methionine and protein N-terminal acetylation were set as variable modifications, with a maximum of 5 variable modifications. Trypsin/P was set as the digestion enzyme, and up to two missed cleavages were permitted. For identification, we applied a maximum false discovery rate of 1% on protein and peptide level. We required 1 or more unique or razor peptides for protein identification. Then, noise was added from the randomly sampled lower range of the limit of detection to all raw intensities. Protein groups with <4.99 average MS/MS counts were removed from further analysis. Protein group intensities were normalized for the total intensity of all observable protein groups in that sample. Normalized protein group intensities were log2-transformed and averaged, from which fold-changes were calculated. P-values were calculated using a two-tailed, heteroscedastic t-test. Gene Ontology analysis was performed using Panther or String. Submitochondrial localization was determined by gene ontology terms for Cellular Component, as well as identified compartment by cross-linked assisted spatial proteomics[1].

### LC-MS/MS analysis total proteome samples

Whole proteome, label-free MS analysis was performed by data-independent acquisition (DIA). For this type of LC-MS/MS analysis, about 1 µg of total peptides were analyzed on a Waters M- Class UPLC using a 15cm IonOpticks Aurora Elite column (75µm inner diameter; 1.7µm particle size; heated to 45°C) coupled to a benchtop Thermo Fisher Scientific Orbitrap Q Exactive HF mass spectrometer. Peptides were separated at a flow rate of 400 nL/min with a 90 min gradient, including sample loading and column equilibration times. Data was acquired in data independent mode using Xcalibur 4.5 software. MS1 Spectra were measured with a resolution of 120,000, an AGC target of 3e6 and a mass range from 350 to 1600 m/z. Per MS1, 29 equally distanced, sequential segments were triggered at a resolution of 30,000, an AGC target of 3e6, a segment width of 43 m/z, and a fixed first mass of 200 m/z. The stepped collision energies were set to 22.5, 25, and 27.

### Quantification and statistical analyses of total proteome samples

All DIA data were analyzed with Spectronaut software version 18.6[3] using directDIA analysis methodology against a UniProt database (Homo sapiens, UP000005640). Carbamidomethylation on cysteines was set as a fixed modification. Oxidation of methionine and protein N-terminal acetylation were set as variable modifications. Trypsin/P was set as the digestion enzyme. Normalization was done per Spectronaut's "automatic normalization". For identification, we applied a maximum false discovery rate of 1% on protein and peptide level. We required 1 or more unique or razor peptides for protein identification. Then, noise was added from the randomly sampled lower range of the limit of detection to all raw intensities. Protein groups with <4.99 average MS/MS counts were removed from further analysis. Protein group intensities were further normalized for the total intensity of all observable protein groups in that sample. Normalized protein group intensities were log2-transformed and averaged, from which fold-changes were calculated. P-values were calculated using a two-tailed, heteroscedastic t-test.

### Purification of SH2 domains

The SHP2 full-length, wild-type gene used as the template for all SHP2 constructs in this study was cloned from the pGEX-4TI SHP2 WT plasmid, which was a generous gift from Ben Neel (Addgene plasmid #8322) (1). SHP2 SH2 domains were cloned into a $His_6$-SUMO-SH2-Avi construct (2). C43(DE3) cells were transformed with plasmids encoding both the respective SH2 domain and the biotin ligase BirA. Cells were grown in LB supplemented with 50 µg/mL kanamycin and 100 µg/mL streptomycin at 37 °C until cells reached an optical density at 600 nm ($OD_{600}$) of 0.5. IPTG (1 mM) and biotin (250 µM) were added to induce protein expression and ensure biotinylation of SH2 domains, respectively. Protein expression was carried out at 18 °C overnight. Cells were centrifuged and subsequently resuspended in lysis buffer (50 mM Tris pH 7.5, 300 mM NaCl, 20 mM imidazole, 10% glycerol, and freshly added 2 mM β-mercaptoethanol). The cells were lysed using sonication (Fisherbrand Sonic Dismembrator), and spun down at 14,000 rpm for 45 minutes. The supernatant was applied to a 5 mL Ni-NTA column (Cytiva). The resin was washed with 10 column volumes lysis buffer and wash buffer (50 mM Tris pH 7.5, 50 mM NaCl, 20 mM imidazole, 10% glycerol, and freshly added 2 mM β-mercaptoethanol). The protein was eluted off the Ni-NTA column in elution buffer (50 mM Tris pH 7.5, 50 mM NaCl, 500 mM imidazole, 10% glycerol) and brought onto a 5mL HiTrap Q Anion exchange column (Cytiva). The column was washed using Anion A buffer (50 mM Tris pH 7.5, 50 mM NaCl, 1 mM TCEP). Protein elution off the column was induced through a salt gradient between Anion A buffer and Anion B buffer (50 mM Tris pH 7.5, 1 M NaCl, 1 mM TCEP). The eluted protein was cleaved at the $His_6$-SUMO tag by addition of 0.05mg/mL $His_6$-tagged Ulp1 protease at 4ºC overnight. This cleavage cocktail was flowed through a 2 mL Ni-NTA gravity column (ThermoFisher) to isolate the cleaved protein away from uncleaved protein and Ulp1. Finally, the cleaved protein was purified by size-exclusion chromatography on a Superdex 75 16/600 gel filtration column (Cytiva) equilibrated with SEC buffer (20 mM HEPES pH 7.4, 150 mM NaCl, and 10% glycerol). Pure fractions were pooled and concentrated, and flash frozen in liquid $N_2$ for long-term storage at −80 °C.

### Purification of full-length SHP2 proteins

Full-length SHP2 variants were cloned into a pET28-His-TEV plasmid from the pGEX-4TI SHP2 WT plasmid. BL21(DE3) cells were transformed with the respective plasmids, and were grown in LB supplemented with 100 μg/mL kanamycin at 37 °C until cells reached an $OD_{600}$ of 0.5. IPTG (1 mM) was added to induce protein expression, which was carried out at 18 °C overnight. Cells were centrifuged and subsequently resuspended in lysis buffer (50 mM Tris pH 7.5, 300 mM NaCl, 20 mM imidazole, 10% glycerol, and freshly added 2 mM β-mercaptoethanol). The cells were lysed using sonication (Fisherbrand Sonic Dismembrator), and spun down at 14,000 rpm for 45 minutes. The supernatant was applied to a 5 mL Ni-NTA column (Cytiva). The resin was washed with 10 column volumes lysis buffer and wash buffer (50 mM Tris pH 7.5, 50 mM NaCl, 20 mM imidazole, 10% glycerol, and freshly added 2 mM β-mercaptoethanol). The protein was eluted off the Ni-NTA column in elution buffer (50 mM Tris pH 7.5, 50 mM NaCl, 500 mM imidazole, 10% glycerol) and brought onto a 5mL HiTrap Q Anion exchange column (Cytiva). The column was washed using Anion A buffer (50 mM Tris pH 7.5, 50 mM NaCl, 1 mM TCEP). Protein elution off the column was induced through a salt gradient between Anion A buffer and Anion B buffer (50 mM Tris pH 7.5, 1 M NaCl, 1 mM TCEP). The eluted protein was cleaved at the $His_6$-TEV tag by addition of 0.10 mg/mL of $His_6$-tagged TEV protease at 4 °C overnight. This cleavage cocktail was flowed through a 2 mL Ni-NTA gravity column (ThermoFisher) to separate the cleaved protein from uncleaved protein and TEV protease. Finally, the cleaved protein was purified by size-exclusion chromatography on a Superdex 200 16/600 gel filtration column (Cytiva) equilibrated with SEC buffer (20 mM HEPES pH 7.5, 150 mM NaCl, and 10% glycerol). Pure fractions were pooled and concentrated, and flash frozen in liquid $N_2$ for long-term storage at −80 °C.

### SH2 specificity profiling

Electrocompetent MC1061 cells were transformed with approximately 100 ng of the strep-tagged $X_5$-Y-$X_5$ library[2]. After 1 hour recovery in 1 mL LB, cells were further diluted into 250 mL LB + 0.1% chloramphenicol. 1.8 mL of overnight culture was used to inoculate 100 mL LB + 0.1% chloramphenicol, and grown until OD600 reached 0.5. 20 mL of cell suspension was induced at 25 °C using a final concentration of 0.4% arabinose until the $OD_{600}$ reached approximately 1 (after about 4 hours). The cells were spun down at 4000 rpm for 15 minutes, and the pellet was resuspended in PBS so that the $OD_{600}$ ~1.5. The cells were stored in the fridge and used within a week.

For each sample, 150 μL of Dynabeads™ FlowComp™ Flexi Kit were washed twice in 1 mL SH2 buffer (50 mM HEPES pH 7.5, 150 mM NaCl, 1 mM TCEP, and 0.2% BSA) on a magnetic rack. The beads were then resuspended in 150 μL of SH2 buffer. SH2 domains were thawed quickly and 20 μM of protein was added to the beads. SH2 buffer was added up to 300 μL, and the suspension was incubated for 1 hour at 4 °C while rotating. After 1 hour, the suspension was placed on a magnetic rack and washed twice with 1 mL SH2 buffer.

150 μL of prepared cells per sample were spun down for 4000 rpm for 5 minutes at 4 °C. Kinase screen buffer was prepared (50 mM Tris, 10 mM magnesium chloride, 150 mM sodium chloride; add 2 mM sodium orthovanadate and 1 mM TCEP fresh) and the cells of each sample were resuspended in 100 μL kinase screen buffer. Kinases c-Src, c-Abl, AncSZ, Eph1B were added to a final concentration of 2.5 μM each, creatine phosphate was added to a final concentration of 5 mM, and phosphokinase was added to a final concentration of 50 μg/mL. The suspension was incubated at 37 °C for 5 minutes before ATP was added to a final concentration of 1 mM. This mixture was incubated at 37 °C for 3 hours. After 3 hours, EDTA was added to a final concentration of 25 mM to quench the reaction. The input library control sample was not phosphorylated. These cells were spun down at 4000 rpm for 15 minutes at 4 °C. The cells were then resuspended in 100 μL of SH2 buffer + 0.1% BSA. Phosphorylation of the cells was confirmed by labeling with the PY20-PerCP-eFluor 710 pan-phosphotyrosine antibody followed by analysis via flow cytometry.

100 μL of phosphorylated cells were mixed with 75 μL SH2-beads for 1 hour at 4 °C while rotating. After 1 hour, samples were placed on a magnetic rack, supernatant was removed and 1 mL SH2 buffer

was added to each sample. This was rotated for 30 minutes at 4 °C to wash the beads. After this wash, the beads were placed on a magnetic rack, the supernatant was removed and 50 µL MilliQ was added.

All SH2-selected samples and the input library control were resuspended in 50 µL MilliQ water, vortexed, and boiled for 10 minutes at 100 °C. The boiled lysate was used as the DNA template in a PCR reaction using the TruSeq-eCPX-Fwd and TruSeq-eCPX-Rev primers. The mixture resulting from this PCR was used directly into a second PCR to append Illumina sequencing adaptors and unique 5' and 3' indices to each sample (D700 and D500 series primers). The resulting PCR mixtures were run on a gel, the band of the expected size was extracted and purified, and its concentration was determined using QuantiFluor® dsDNA System (Promega). Samples were pooled at equal molar ratios and sequenced by paired-end Illumina sequencing on a MiSeq or NextSeq instrument using a 150 cycle kit. The number of samples per run, and the loading density on the sequencing chip, were adjusted to obtain at least 1-2 million reads for each index/sample.

### Analysis of data from screens with the $X_5$-Y-$X_5$ Library

Deep sequencing data were processed and analyzed as described previously[2]. First, paired-end reads were merged using FLASH[3]. Then, adapter sequences and any constant regions of the library flanking the variable peptide-coding region were removed using Cutadapt[4]. Finally, these trimmed files were analyze using in-house Python scripts in order to count the abundance of each peptide in the library, as described previously (https://github.com/nshahlab/2022_Li-et-al_peptide-display). For each individual amino acid, we counted its occurrence at every position along peptides of the expected length of 11 residues, excluding any sequences containing a stop codon. This generated an 11x20 counts matrix with each position in the peptide represented by a column (from –5 to +5), and each row represented by an amino acid (ordered by biochemical properties). Frequencies of each amino acid at each position were determined by taking the position-specific count for each amino acid and dividing that by the column total. Frequencies in a matrix from a selected sample were further normalized against frequencies from an input sample, and the resulting enrichment values were $\log_2$-transformed. Matrices from two independent screens with each SH2 domain were averaged to yield the data **Figure 2E** and **Supplementary Table 4**.

### Analysis of human phosphosites using $X_5$-Y-$X_5$ library screens

39235 human phosphosites were downloaded from PhosphoSitePlus (retrieved on October 20, 2022). In order to score each phosphosite using the generated position-weighted counts matrices from our $X_5$-Y-$X_5$ library screen, we first calculated the normalized enrichment for each amino acid at each position across the matrices. Then, for each phosphosite, we summed up the $\log_2$-normalized enrichments for each residue according to the enrichment matrix (excluding the central tyrosine), and divided the sum by the number of scored residues (10 in total). Finally, the scores for each screen were normalized such that the minimum and maximum possible theoretical score was set to 0 and 1, respectively. For each SH2 domain, two replicate screens were conducted, and the scoring matrices from each individual replicate were used to score all phosphosites. Then, the scores from each replicate were averaged to yield the data in **Supplementary Table 4**.

### DNA constructs

The SHP2 gene was cloned from the pGEX-4TI SHP2 WT plasmid from Ben Neel (Addgene plasmid #8322). TurboID was cloned from the V5-TurboID-NES_pCDNA3 plasmid, which was a gift from Alice Ting (Addgene, #107169). The mouse c-Src gene was expressed from the pCMV5 mouse Src plasmid, a gift from Joan Brugge and Peter Howley (Addgene plasmid #13663). We received the R777-E227 Hs.RASA1 plasmid as a gift from Dominic Esposito (Addgene, # 70511). The pCDNA3-hPZR-WT plasmid was a gift from Anton Bennett.

### Purification of SHP2-TurboID constructs

BL21 (DE3) cells were transformed with $SHP2^{WT}$-TurboID or $SHP2^{E76K}$-TurboID DNA. After heat shock, cells were recovered in 1 mL Luria Broth (LB) for 1 hour at 37 °C while shaking. 100 µL was plated and incubated overnight at 37 °C. The next morning, colonies were resuspended and grown in LB with 100 µg/mL kanamycin at 37 °C until the culture reached an $OD_{600}$ of 0.5. 1 mM Isopropyl β-d-1-thiogalactopyranoside (IPTG) was added to induce protein expression, which proceeded at 18 °C overnight. Cells were pelleted and resuspended in lysis buffer (50 mM Tris pH 7.5, 300 mM NaCl, 20 mM imidazole, 10% glycerol, and freshly added 2 mM β-mercaptoethanol). The cells were lysed by sonication (Fisherbrand Sonic Dismembrator), and centrifuged at 14,000 rpm for 45 minutes. The supernatant was brought onto a 5 mL Ni-NTA column (Cytiva). The column was washed with 10 column volumes lysis buffer and wash buffer (50 mM Tris pH 7.5, 50 mM NaCl, 20 mM imidazole, 10% glycerol, and freshly added 2 mM β-mercaptoethanol). The protein was eluted off the Ni-NTA column in elution buffer (50 mM Tris pH 7.5, 50 mM NaCl, 500 mM imidazole, 10% glycerol) and applied to a 5mL HiTrap Q Anion exchange column (Cytiva). The column was then washed using Anion A buffer (50 mM Tris pH 7.5, 50 mM NaCl, 1 mM tris(2-carboxyethyl)phosphine (TCEP)). The protein was eluted off the column through a salt gradient between Anion A buffer and Anion B buffer (50 mM Tris pH 7.5, 1 M NaCl, 1 mM TCEP). The eluted protein was cleaved at the His6-TEV tag by addition of 0.10 mg/mL of His6-tagged TEV protease at 4 °C overnight. This cleavage mixture was applied to a 2 mL Ni-NTA gravity column (ThermoFisher) to separate the cleaved protein from uncleaved protein and TEV protease. Finally, the cleaved protein was purified by size-exclusion chromatography on a Superdex 200 10/300 gel filtration column (Cytiva) equilibrated with SEC buffer (20 mM HEPES pH 7.5, 150 mM NaCl, and 10% glycerol). Pure fractions were pooled and concentrated, and flash frozen in liquid N2 for long-term storage at −80 °C.

### Peptide activation assay

Full-length SHP2 and SHP2-TurboID constructs were diluted in assay buffer (60 mM HEPES pH 7.2, 75 mM KCl, 75 mM NaCl,1 mM EDTA, 0.05% Tween-20, with 0.5 mM TCEP freshly added) to a 2X concentration of 0.1 nM. Peptides were serially diluted in assay buffer, with the last point of the concentration series lacking peptide. 15 µL of peptide dilution mix and 15 µL of 800 µM DiFMUP was added to a black 96-well half area plate. 30 µL SHP2 was added to each well to a final concentration of 0.05nM immediately before starting the plate reader assay. Kinetic measurements (Ex 358/Em 355) were taken on a BioTek Neo2 plate reader every 25 seconds for 6 minutes.

### Co-immunoprecipitation experiments

1 x $10^6$ SHP2 KO HEK 293 cells were seeded in a 6 cm plate. The next day, cells were transfected using 2 µg of each plasmid (SHP2 interacting protein of interest; for MPZL1 IPs c-Src was co-transfected as well), and 18 µg PEI in 600 µL DMEM. The transfection medium was refreshed after 16 hours and replaced with complete medium. After 48 hours, cells were harvested by scraping in PBS. Cells were washed 3 times in 1 mL PBS, and lysed in 100 µL lysis buffer (20 mM Tris-HCl, pH 8.0, 137 mM NaCl, 2 mM EDTA, 10% glycerol, and 0.5% NP-40 + protease inhibitors + phosphatase inhibitors) for 30 minutes while rotating at 4 °C. Cells were spun at 17.7 rpm for 15 minutes at 4 °C. Supernatant was transferred to a clean Eppendorf tube and stored at -20 °C.

Protein concentration was determined using a bicinchoninic acid (BCA) assay and absorbance was measured at 562 nm using a BioTek Synergy Neo2 multi-mode reader. 300 µg of protein in a total volume of 380 µL was incubated overnight with 30 µL of magnetic Myc-beads while rotating at 4 °C overnight. The beads were washed 3 times on a magnetic rack using 1 mL lysis buffer. Then, 65 uL1x Laemmli buffer was added and beads were boiled at 100ºC for 8 minutes. For whole cell lysates, 15 µg protein was loaded onto a gel. For IP samples, 15 µL of boiled supernatant was used. Gel was transferred onto a nitrocellulose membrane using TurboBlot (BioRad) and the membrane was blocked using 5% bovine serum albumin (BSA) in Tris-buffered saline (TBS) for 1 hour at room temperature. Membranes were rinsed with TBS with 0.1% Tween-20 (TBS-T) and incubated with primary antibodies in TBST + 5% BSA overnight at 4ºC (Src 1:1000, β-actin 1:5000, Myc 1:5000, FLAG 1:5000, PD-1 1:1000, pTyr 1:2000). Co-immunoprecipitation of the protein of interest was detected using an α-FLAG antibody. Membranes were washed and incubated with secondary antibodies (IRDye 680 and 800). Blots were imaged on a LiCor Odyssey. Band intensities were quantified using Image Studio Lite (Version 5.2) using the Median Background setting. IP intensities were divided by corresponding intensities in total cell lysate. P-values were calculated in GraphPad Prism (Version 10.1.0) using a paired, one-sided T-test.

### Ras dephosphorylation assay (Rassay)

0.8 x $10^6$ $SHP2^{KO}$ HEK 293 cells were seeded in a 6 cm plate. The next day, cells were transfected with Ras, Ras and Src, or Ras, Src and SHP2; to a total of 3 µg in 300 µL DMEM with 9 µg of polyethyleneimine. The transfection medium was refreshed the next morning and replaced with warm DMEM with 10% FBS. Approximately 36 hours after transfection, the cells were harvested by scraping and washed 3 times in 1 mL cold PBS. Cells were lysed in 150 µL lysis buffer (20 mM Tris pH 8.0, 137 mM NaCl, 2 mM EDTA, 10% glycerol, 0.5% NP-40, with freshly added phosphatase- and protease inhibitors) for 25 minutes on ice. Lysates were spun down for 15 minutes at 17,000 g in a 4 °C tabletop centrifuge. Supernatant was used in a bicinchoninic acid (BCA) assay to determine protein concentration. 80 µg of protein was used in an immuno-precipitation (IP) with 5 µg of packed Pierce anti-HA magnetic beads (Fisher, #88836) in a total volume of 350 µL lysis buffer. Samples were incubated at 4 °C overnight while rotating. The next morning, beads were washed 3 times on a magnetic racks using 1 mL of lysis buffer, and resuspended in 65 µL of 1x Laemmli buffer. All samples were boiled at 100 °C for 8 minutes, and 15 µg of total protein (total cell lysate) or 15 µL of each sample (immuno-precipitation) was loaded onto a 12% acrylamide gel. Proteins were transferred to a 0.45 µm nitrocellulose membrane using the StandardSD protocol on the Bio-Rad Trans-Blot Turbo. Membranes were blocked for 1 hour at room temperature using 5% BSA in TBS. Primary antibodies were stained for 2 hours at room temperature (MPZL1 TCL: Src 1:1000, β-actin 1:5000, Myc 1:5000, HA 1:1000; MPZL1 IP: HA 1:1000, pTyr 1:2000; ATPAF1 TCL: Vinculin 1:1000, FLAG 1:5000, Myc 1:5000; ATPAF1 IP: FLAG 1:5000, Myc 1:5000) in 5% BSA in TBST. Membranes were washed 3 times in 5 mL TBST for 5 minutes each. Secondary antibodies were incubated in 5% BSA in TBST for 1 hour at room temperature (1:10,000). Membranes were imaged on a LiCor Odyssey and bands were quantified using ImageStudio.

| **Description** | **Vendor** | **Catalog #** |
|---|---|---|
| Myc (Clone (9E10) | Invitrogen | R95025 |
| c-Src | CST | 2123S |
| B-actin | Sigma | A5441 |
| Phospho-Tyrosine (P-Tyr-1000) MultiMab | CST | 8954S |
| Pzr | CST | 9893S |
| FLAG | MPBio | 08L100031 |
| Vinculin | CST | 13901S |
| HA (TCL) | Sigma | SAB5600116 |
| HA (IP) | Sigma | 11867423001 |
| IRDye® 680RD Goat anti-Rabbit IgG | LiCor | 926-68071 |
| IRDye® 800CW Goat anti-Mouse IgG | LiCor | 926-32210 |
| IRDye 800CW Goat anti-Rat IgG | LiCor | 926-32219 |

### Fluorescence microscopy

$10^6$ U2-OS cells were seeded in a black, glass bottom 6-well plate. The next day, cells were transfected with 1 µg of SHP2 DNA in empty DMEM overnight. Medium was refreshed with DMEM + 10% FBS. 48 hours after transfection, half the media was replaced with 4% paraformaldehyde (PFA) + 4% sucrose in PBS, pre-warmed to 37 °C. This was incubated for 1 minute, then the entire well-volume was replaced with 4% PFA + 4% sucrose in PBS. The plate was incubated for 10 minutes at 37˚C. Wells were rinsed twice with 1mL PBS and permeabilized with 0.1% TritonX-100 in PBS for 10min on ice. Wells were rinsed twice with 1mL PBS after which primary antibodies were applied in PBS + 0.05% Tween-20 (α-Tomm20, Sigma, # ZMS1118, 1:100; and α-SHP2, Cell Signaling Technology (CST), # D50F2, 1:100) for 1 hour at room temperature. After incubation, wells were rinsed 3 times with 1 mL PBS. Secondary antibodies were applied in PBS (Anti-rabbit IgG (H+L), F(ab')2 Fragment (Alexa Fluor® 488 Conjugate), CST, # 4412, 1:2000; Goat Anti-Mouse, Alexa555, ThermoFisher Scientific, A-21424, 1:1000), Alexa Fluor® 647 Phalloidin, CST, #8940, 1:100) for 30 minutes at room temperature in the dark. Wells were rinsed 3 times with 1 mL PBS. 1 µg/mL 4′,6-diamidino-2-phenylindole (DAPI) was added in PBS. All images were acquired under oil immersion 60x magnification (Nikon, MRD71670) using a confocal spinning disk microscope (Andor Dragonfly) coupled to a Nikon Ti-2 inverted epifluorescence microscope with automated stage control, Nikon Perfect Focus System and a Zyla PLUS 4.2-megapixel USB3 camera. Illumination was done with 100 mW 405 nm, 50 mW 488 nm, 50 mW 561 nm and 140 mW 640 nm solid-state lasers. All hardware was controlled using Andor Fusion software. Lasers, laser powers, exposure times, objectives and experiment-specific acquisition parameters are 100% power 100ms exposure for all the images. Images were acquired with 11 z-slices at 2.0-µm intervals (Total scan size 20 µ m). Images were analyzed in ImageJ/Fiji. The z-slice with the clearest mitochondrial signal (Tom20) was selected, and pixels were selected that had signal for both SHP2 and Tom20.

### Differential scanning fluorimetry

Purified protein stocks were thawed and diluted in DSF buffer (20 mM HEPES pH 7.5, 50 mM NaCl, 0.4% DMSO). 19 µL of buffer was added to a MicroAmp Fast Optical 96-well Reaction plate (Applied Biosystems, # 4346906). 1 µL of 500x SYPRO Orange Protein Gel Stain (Thermo Fisher, catalog no. S-6650) was added to a final protein concentration of 10 µM and 25x SYPRO Orange. Melting curves were performed in an Applied Biosystems Step-One Plus RT-PCR thermocycler. Temperature measurements started at 15 ºC, and temperature was raised by 0.5 ºC every minute with continuous measurements of fluorescence (excitation: 472 nm; emission: 570 nm). Raw fluorescence values along with corresponding temperatures were analyzed using DSFworld and $T_m$ values were calculated using dRFU.

### Analysis of solvent exposure in molecular dynamics simulations

Molecular dynamics simulations were performed and reported previously[5]. In total, twelve 2.5 µs simulations were analyzed: 3 simulations of $SHP2^{WT}$ and 3 simulations of $SHP2^{E76K}$, starting from the conformation seen in PDB code 4DGP, and 3 simulations of $SHP2^{WT}$ and 3 simulations of SHP2E76K, starting from the conformation seen in PDB code 6CRF. MD trajectories were compiled from the raw data using the CPPTRAJ module of AmberTools22[6]. Structures were extracted from the trajectories 10 ns increments for analysis. Solvent-accessible surface area (SASA) of each residue was calculated at each extracted time point using the PDB module in Biopython[7]. SASA values for all simulations starting from 4DGP or 6CRF were combined to generate the distributions shown in **Figure 6D**.

### Mitochondrial isolations

10 x $10^6$ $SHP2^{KO}$ HEK 293 cells were seeded in a 15 cm plate. Cells were transfected with 25µg DNA in DMEM with 75 µg of PEI. Transfection medium was aspirated and replaced with DMEM + 10% FBS. 36 hours after transfection, cells were harvested by scraping and washed 3 times in 1 mL PBS. Cell pellets were processed using a Mitochondria/Cytosol Fractionation Kit (Fisher, #89874) according to manufacturer's instructions. Mitochondrial pellet was lysed in 30-50 µL lysis buffer (20 mM Tris HCl pH 8.0, 137 mM NaCl, 10% glycerol, 1% Nonidet P-40 (NP-40), 2 mM EDTA, with protease inhibitors and phosphatase inhibitors added fresh). Protein concentration of cytosolic and mitochondrial fractions was determined using a bicinchoninic acid (BCA) assay and samples were further analyzed by Western blot.

Quantification of reactive oxygen species

Reactive oxygen species (ROS) were profiled using MitoSOX™ Mitochondrial Superoxide Indicators for live-cell imaging (Invitrogen, # M36006) according to manufacturer's instructions. Briefly, 0.8 x $10^6$ SHP2$^{KO}$ HEK 293 cells were seeded in a 6 cm plate, and transfected the next day with 5 µg of SHP2 DNA in empty DMEM. Transfection medium was replaced the following morning with DMEM + 10% fetal bovine serum (FBS). Cells were harvested 36 hours post-transfection by scraping in PBS, and washed 2 more times in 1 mL PBS. MitoSox Green reagent was diluted to 1mM in dimethylformamide (DMF), and further diluted to a working concentration of 2.5 µM in PBS. Cells were resuspended in 200 µL of MitoSox Green working solution for 1 hour at 37 °C; + ATP samples were incubated with 5mM ATP and MitoSox Green simultaneously. Cells were washed 3 times in 500 µL PBS and analyzed on an Attune NxT using 488 nm excitation and emission filter 530/30.
